# Functional Anatomy, Biomechanical Performance Capabilities and Potential Niche of StW 573: an *Australopithecus* Skeleton (circa 3.67 Ma) From Sterkfontein Member 2, and its significance for The Last Common Ancestor of the African Apes and for Hominin Origins

**DOI:** 10.1101/481556

**Authors:** Robin Huw Crompton, Juliet McClymont, Susannah Thorpe, William Sellers, Jason Heaton, Travis Rayne Pickering, Todd Pataky, Dominic Stratford, Kristian Carlson, Tea Jashashvili, Amélie Beaudet, Laurent Bruxelles, Colleen Goh, Kathleen Kuman, Ronald Clarke

## Abstract

StW 573, from Sterkfontein Member 2, dated ca 3.67 Ma, is by far the most complete skeleton of an australopith to date. Joint morphology is in many cases closely matched in available elements of *Australopithecus anamensis* (*eg.* proximal and distal tibial and humeral joint-surfaces) and there are also close similarities to features of the scapula, in particular, of KSD-VP-1/1 *A. afarensis* from Woranso-Mille. The closest similarities are, however, to the partial skeleton of StW 431 from Sterkfontein Member 4. When considered together, both StW 573 and StW 431 express an hip joint morphology quite distinct from that of *A. africanus* Sts14, and a proximal femur of a presumed *A. africanus* from Jacovec Cavern at Sterkfontein, StW 598. This, and other evidence presented herein, suggests there are two pelvic girdle morphs at Sterkfontein, supporting Clarke (2013) in his recognition of a second species, *A. prometheus,* containing StW 573 and StW 431. StW 573 is the first hominid skeleton where limb proportions are known unequivocally. It demonstrates that some early hominins, at the time of formation of the Laetoli footprints (3.6 Ma), were large-bodied. with hindlimbs longer than forelimbs. Modelling studies on extant primates indicate that the intermembral index (IMI) of StW 573, low for a non-human great ape, would have substantially enhanced economy of bipedal walking over medium-to-long distances, but that it was still too high for effective walking while load-carrying. It would, however, have somewhat reduced the economy of horizontal climbing, but made *Gorilla-*like embracing of large tree-trunks less possible. Consideration of both ethnographic evidence from modern indigenous arboreal foragers and modern degeneracy theory cautions against prescriptive interpretations of hand- and foot-function, by confirming that both human-like upright bipedalism and functional capabilities of the hand and foot can be effective in short-distance arboreal locomotion.

## 1. Introduction

While it is now largely accepted that there was no phase of terrestrial knucklewalking in hominin evolution (see eg., Dainton and Macho, 1999; Dainton, 2001; Clarke, 2002; Kivell and Schmitt, 2009), and that australopiths show adaptations to both terrestrial bipedalism and arboreal locomotion, there is still no firm consensus on whether the ‘arboreal’ features of australopith postcrania would have been the subject of positive selection or were selectively neutral anachronisms (Ward 2002, 2013). The view that the two activities must be substantially mechanically incompatible is still current (see eg., Kappelman et al., 2016). The two alternative paradigms date from extended debates (see eg., Latimer, 1991 versus Stern and Susman, 1991) concerning the significance of the AL-288-1 ‘Lucy’ skeleton of *Australopithecus afarensis* in 1974. Although some one-third complete, this partial skeleton, and other more recently discovered partial australopith skeletons (eg., the Woranso-Mille *Australopithecus afarensis* skeleton, KSD-VP-1/1 ca. 3.6 Ma [Haile-Selassie et al. 2010]) and the Malapa *A. sediba* skeletons, MH-1 and MH-2, ca. 1.977 Ma [Berger, 2013]) are too incomplete to provide reliable upper and lower limb lengths, a crucial variable in assessing terrestrial and arboreal locomotor performance capabilities, and lack other auxiliary features which might offer a clear signal of the existence of selection for arboreal performance capabilities.

It has become increasingly clear, since the discovery of the foot bones of StW 573 (‘Little Foot’) some 23 years ago (Clarke and Tobias, 1995), that the continued painstaking freeing of Little Foot’s fragile bones from their matrix of hard breccia might for the first time provide unequivocal data on australopith postcranial anatomy. Some missing small skeletal elements may be recovered with further excavation, and some (such as the distal foot) were destroyed by lime mining a very long time ago, but at least 90% of the skeleton, complete enough for unequivocal knowledge of limb lengths, and with remarkably good preservation of joint surfaces and other vital detail has now been excavated and prepared (Clarke 2018, Figure 26). This completeness is unmatched until *Homo ergaster* KNM-WT 15000, at 1.5 Ma. After much debate (see Bruxelles et al. 2018, submitted), the age of this specimen is now confidently set at ca 3.67 Ma, very close to the age of the Laetoli footprint trails (Leakey and Hay, 1979), which were previously our best source of information on the locomotor capabilities of australopiths. Now, StW 573 for the first time offers unequivocal information on limb proportions of forelimb and hindlimb.

Here we review aspects of contributions on StW 573 which bear on its potential niche as an individual. These aspects include the geological and palaeoenvironmental context (Bruxelles et al. 2018, submitted), its craniodental anatomy (Clarke and Kuman, 2018, submitted), the endocast (Beaudet et al., 2018a, in press), the inner ear (Beaudet et al. 2018b, submitted), the scapula and clavicle (Carlson et al., 2018, in prep.), the hand (Jashashvili et al., 2018 in prep.) and the foot (Deloison, 2004). Heaton et al. (2018, submitted) focus on statistical/morphometric descriptions of the longbones of StW 573 and comparisons with other species, and this paper is referred to where appropriate to avoid duplication. However, the present paper is quite distinct in its focus on: a) comparisons of those aspects of joint shape which in the literature are generally regarded as informative concerning locomotor function, and b) reconstruction of StW 573’s potential ecological niche by proxy experiments and extant in-silico modelling. We adopt the hypothesis-based approach of Wainwright (1991) to species ecomorphology, which focuses on individual performance (and in this respect differs considerably from Bock and Von Wahlert’s (eg. 1965) previous formulation which includes form, function and biological role but not performance, and is not therefore suited to experimental testing). This paper, and Heaton et al. (2018, submitted) are thus distinct in aims and methods, and while they refer to each other where appropriate and may indeed to be read in tandem as complementary/companion papers, they can equally stand, and can be assessed, quite independently. In addition, we pay close attention to the significance of modern ecological dynamics theory, particularly plasticity (see eg. Neufuss et al. 2014); and neurobiological degeneracy (see eg. Edelman and Gally, 2001), for the cheiridia (hands and feet) in particular, which suggest that circumspection needs to be applied to functional interpretation of foot (see eg. Deloison, 2004) and hand morphology.

Further, euhominoids (otherwise known as crown hominoids, so excluding eg. Proconsulidae, Pliopithecidae etc) display high levels of plasticity in muscle architecture: we have already noted that while DeSilva (2009) asserted that humans could not achieve the required dorsiflexion for chimpanzee-like vertical climbing, Venkataraman et al. (2013) showed that (presumably developmental) fibre-length plasticity enables some forest hunter-gatherers to do so, while neighbouring non-climbing populations cannot. Further, Neufuss et al. (2014) showed that while lemurs, like all primarily pronograde mammals studied to date, exhibit a dichotomy in axial musculature between deep slow contracting local stabilizer muscles and superficial fast contracting global mobilizers and stabilizers, hominoids, as previously shown for *Homo*, show no regionalization. Thus, it appears that hominoids have been under selective pressure to develop and sustain high functional versatility of the axial musculature, reflecting a wide range of mechanical demands on the trunk in orthogrady. Neufuss et al. (2014). Using this analytical framework, focusing throughout on StW 573’s individual locomotor performance capabilities, we hope to advance understanding of her biomechanical interaction with her environment and potential niche: her ecomorphology sensu Wainwright (1991).

## Background

### 2.1. The Sterkfontein Formation

The Sterkfontein Formation is the world’s longest succession documenting the evolution of hominins, fauna, landscape and ecology. Member 5 holds *Homo habilis* and *Homo ergaster*, *Paranthropus,* and evidence for over half a million years’ evolution of early stone tool technologies. Member 4, dating to between at least 2.8 Ma. and 2.15 Ma, is the richest *Australopithecus*-bearing deposit in the world and contains two species of *Australopithecus*, together with diverse and extensive faunal evidence. Four metres below, under the yet unexplored and extensive Member 3, which potentially documents a long period of evolution of *Australopithecus,* lies Member 2, which has yielded the world’s most complete *Australopithecus* skeleton, StW 573, the only hominin found in this deposit. Member 2 formed around 3.67 Ma (Granger et al. 2015) based on isochron burial dating using cosmogenic aluminium-26 and beryllium-10. The detailed descriptions of the Member 2 sedimentary units by Bruxelles et al. (2014, 2018 submitted) confirm the 3.67 Ma age of the skeleton published by Granger et al. (2015). Pickering and colleagues’ recent (2018) repetition of a 2.8 Ma date fails to take into account, and fails to cite, Bruxelles and colleagues’ (2014) demonstration that the flowstones are intrusive and dates for StW 573 based on them thus invalid.

Importantly, taphonomic evidence (Clarke 2018 submitted; Bruxelles et al. 2018 submitted) indicates that StW 573 died and was fossilized (below the ecological context in which she lived), from a fall into a steep cave shaft leading to an underground cavern. The skeleton is associated, in Member 2, with fauna dominated by cercopithecoids and carnivores (see section 2.4).

### 2.2 The StW 573 partial skeleton

The skeletal elements found to date are shown in assembly in Figure 26 of Clarke (2018, submitted). The skeleton offers, for the first time in one individual *Australopithecus*, complete (if deformed) skull and mandible, many vertebrae and ribs, a crushed pelvis and ischiopubic ramus, femora (broken but with overlapping morphology allowing confident length reconstruction), one intact and one slightly damaged but measurable tibia, partial left and right fibulae which overlap sufficiently to be sure of length and morphology, a partial foot (representing primarily the medial column) foot, two scapulae (one articulated with the upper limb), both claviculae (one partial and one complete), both humeri (one partially crushed), both radii and ulnae (one side near-intact and the other crushed and deformed most probably by a badly healed injury in-vivo), and finally one partial and one virtually complete hand (missing only one distal phalanx).

Their taphonomy and condition are discussed in detail in Clarke (2018, submitted) and the stratigraphic context in Bruxelles et al. (2018, submitted). StW 573’s pelvis was substantially flattened post-mortem, but preservation of its margins is good enough to identify an obtuse greater sciatic notch angle (Figure 1 top) and hence suggest female sex. In contrast Figure 1 bottom shows the original reconstruction of the pelvis of StW 431 from Member 4, which appears closely similar but is indicative of a male from the acuteness of the greater sciatic notch. Lipping of the margins of the vertebral bodies of StW 573 (Figure 1, top) and heavy toothwear (Clarke et al., 2018, submitted), indicate that she was an old individual. StW 573 would have been some 130 cm in stature (RJC pers.comm. to RHC), which is some 10 cm. less than the average for modern Bolivian women, the world’s shortest female population. By contrast, the stature of AL-288-1 would have been some 107 cm (Jungers, 1988). The considerable difference in stature is in accord with conclusions from the dimensions of the penecontemporaneous Laetoli footprints (both Deloison [1993, pp. 624-629], for Laetoli G and Masao et al. [2016] for the more extensive Laetoli S) that there was a large range in stature in early hominins. The slightly younger KSD-VP-1/1 partial skeleton confirms this conclusion for *A. afarensis*, to which it is referred.

**Figure 1.**
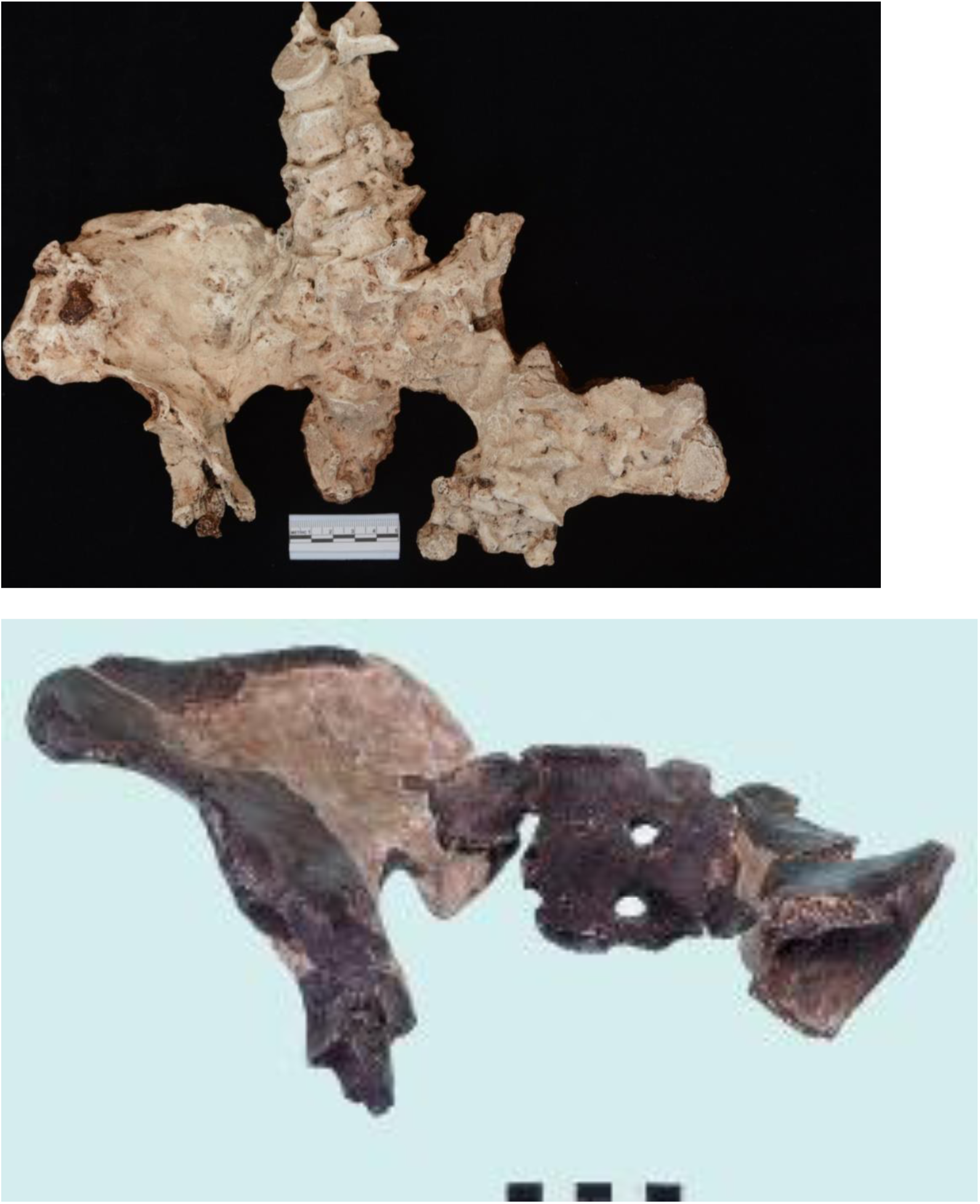
Top: Pelvis of StW 573, Bottom: the Robinson (1972) reconstruction of the pelvis of StW 431 (archival photograph)

### 2.3. Environmental Context

While based on the bovids, Vrba (1975) suggested a medium density woodland with a substantial open component, and based on the overall community of mammals, Reed (1997) similarly suggested open woodland with bush, However, most of the Sterkfontein Member 2 fauna represent cercopithecoid or carnivore taxa which are today habitual climbers, and ancient gravels indicate a large, slow flowing river in the base of the valley (Pickering et al., 2004). Similarly, Elton et al. (2016) indicate that the cercopithecoids which are found in Member 2 were probably to some extent ecologically dependent upon trees for foraging, predator avoidance, or both. Thus, Pickering et al., (2004).suggest a paleohabitat of rocky hills covered in brush and scrub, but valley bottoms with riverine forest, swamp and standing water. Such a paleoenvironment might resemble that in today’s Odzala-Koukoua National Park, Congo, where grassland, standing water and forest are interspersed (see eg., https://reefandrainforest.co.uk/news-item/trip-report-wildlife-republic-congo. Member 4 preserves a takin-like (and hence presumably also woodland) bovid *Makapania*, as well as large cercopithecoids (Pickering et al. 2004), which are associated with forest vines requiring large trees (Bamford, 1999), including one today known exclusively from central and Western African tropical forest. This evidence suggests that little dessication occurred until Member 5 times. A carbon isotope study of faunal teeth by Luyt and Lee Thorpe (2003) also confirms that a drier, more open environment was only established by 1.7 Ma at Sterkfontein.

### 2.4. Species affinities

There has been a long debate concerning the number of species within *Australopithecus* in South Africa, and particularly with relationship to the validity of *Australopithecus prometheus*. Grine (2013) feels that craniodental and some ancillary paleoenvironmental data are insufficient to justify splitting *A. africanus*, but Clarke (2013) presents craniodental and also postcranial evidence for species diversity at Sterkfontein. Of particular relevance here, he finds a distinction between *A. prometheus*, represented by, for example, the partial skeleton of StW 431 and the near-complete StW 573 skeleton and *A. africanus,* represented by Sts 14. Most of these discussions have focused on dental and gnathocranial evidence, but recent postcranial discoveries of postcrania have broadened the evidence base considerably, with particular attention now being paid to the pelvis and hip joint, which are crucial to both obstetric and locomotor evolution.

Thus, Figure 2 shows that the ilia of the Member 4 StW 431 and the Member 2 StW 573 are closely similar in size and shape, even though the greater sciatic notches indicate that StW 573, with an obtuse notch, is most likely a female, and StW 431, with an acute notch, is apparently a male. By contrast, Figure 3 shows that the ilia of Sts14 and *A. afarensis* AL-288-1 -- from their obtuse greater sciatic notches both apparent females – are substantially smaller. While Toussaint et al. (2003), who described StW 431, refer it to *A. africanus*, co-author Macho prefers attribution to *A. prometheus* (G.A.M. pers. comm. to RHC). Importantly, both the unreconstructed pelvis described by Toussaint et al. (2003) and the Kibii and Clarke (2003) reconstruction show that the acetabular margin is well preserved, and acetabular size well defined. Toussaint et al. (2003) note: ‘The acetabulum is clearly large: to judge by the preserved part, its vertical diameter would exceed 42 mm compared with 29.2 mm in Sts 14’ (page 219).

**Figure 2.**
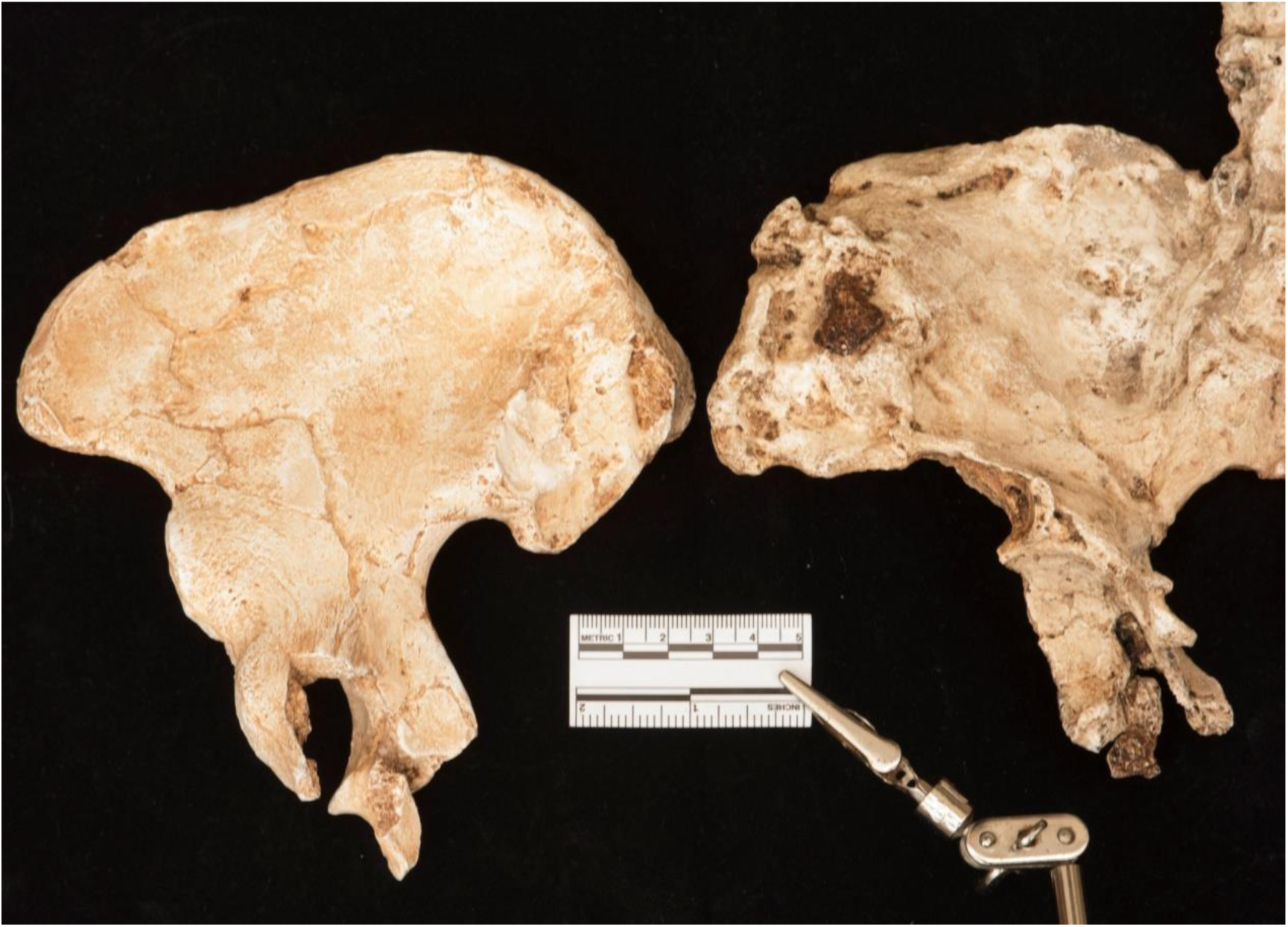
Innominates of (Left) StW 431 (Right) StW 573

**Figure 3.**
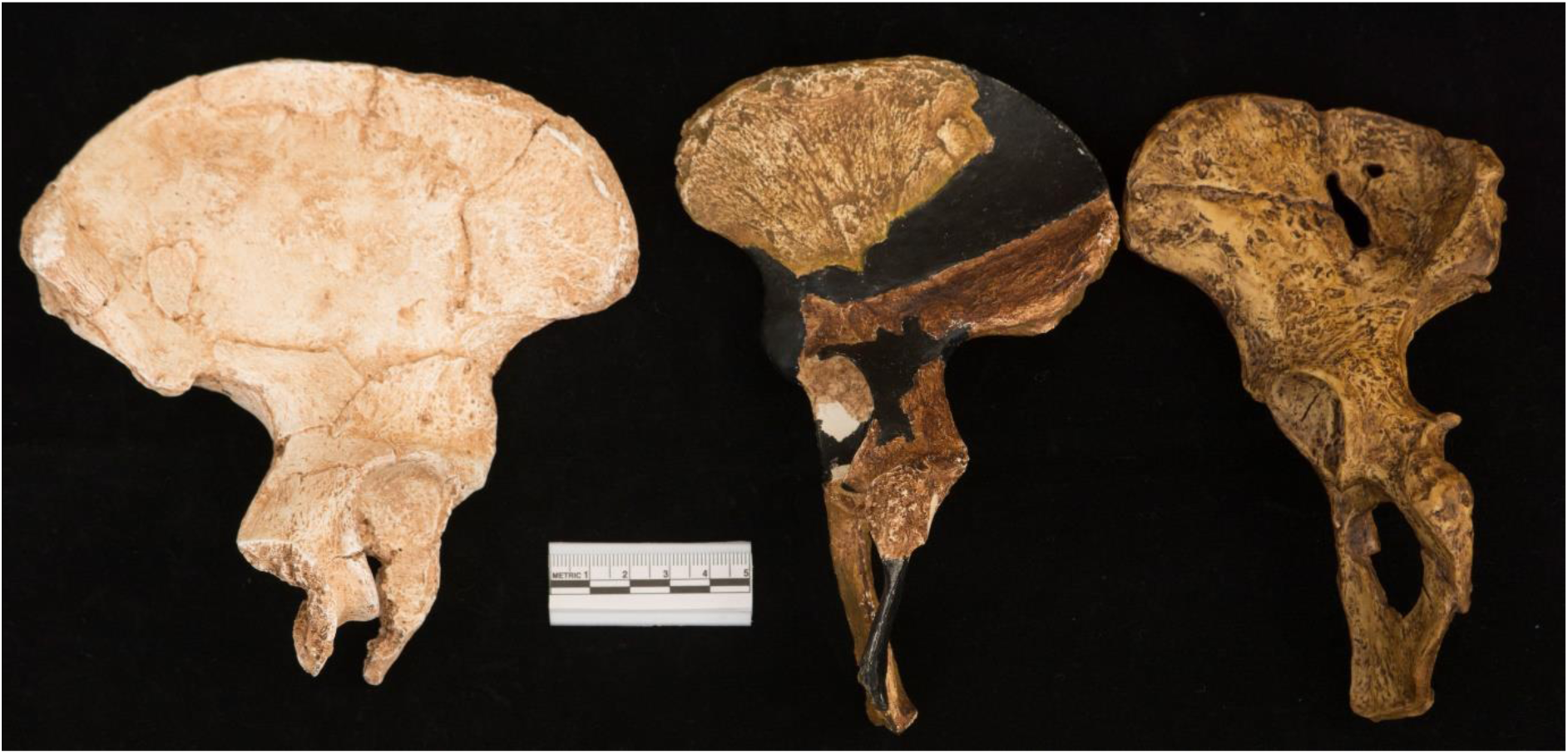
Innominates of (Left to Right) StW 431, Sts 14 and AL-288-1

Acetabular size is also large in StW 573, probably over 36 mm (Figure 4), although the acetabulum has been compressed by taphonomic events. Figures 5 and 6 show that not only is the acetabulum large, but the StW 573 femoral head is a close match for the acetabulum of StW 431, while the head of the *A. africanus* proximal femur from Jacovec Cavern StW 598 is markedly smaller than the acetabulum of StW 431 or StW 573. The Jacovec Cavern proximal femur (StW 598) is, however, a good match for the acetabulum of Sts 14 (Figure 7 top left). That this reflects more than allometry is shown by the fact that the femur of *A. afarensis* also has a small head but lacks an obviously long femoral neck. We predict that other isolated material referable to *A. africanus* will similarly be found to have a long femoral neck. The StW 367 femur from Member 4 Sterkfontein shows a remarkable similarity to that of Jacovec StW 598 (Figure 7, top right). Thus both the small-bodied, long femoral neck/small femoral head morph (eg. Jacovec, StW 367) and the large-bodied, short femoral neck but large femoral head large hip joint morph (StW 431, StW 573) were present in both Member 2 and Member 4 times. The StW 573 femur resembles those of both humans and KNM WT 15000 (Figure 7, bottom) (and see Heaton et al. 2018, submitted, for morphometric detail). A large femoral head is commonly, and reasonably, associated with large forces operating across the hip joint and may be expected to correlate with body size. Femoral neck length, however, is likely related to the moment arm of the hip abductors (see eg. McHenry, 1975), but in what way, and with what iliac geometries, remains to be tested. It should be noted that a long femoral neck can prima facie be assumed to increase the risk of femoral neck fracture during instability events or falls, as it will increase the moment arm about the femoral neck from the impact or instability site. Stern (2000) is cited by Toussaint *et al.* (2003) in reference to the possibility that *A. afarensis* and *A. africanus* may have had a less effective abduction capacity in gluteus medius ‘thus compromising stabilization during walking’ (page 222). This could readily, and will be, tested in silico.

**Figure 4.**
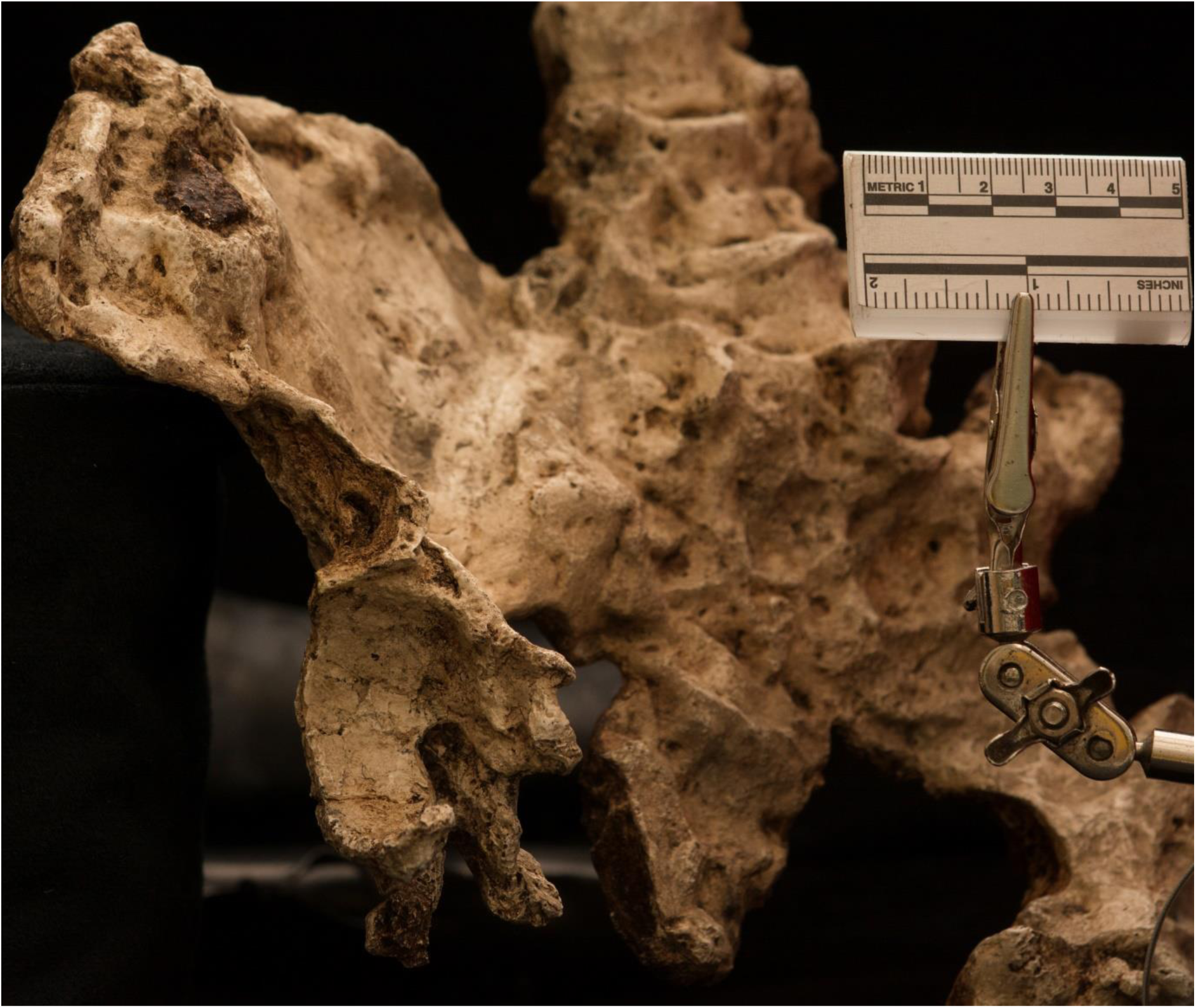
Innominate of StW 573 showing acetabulum

**Figure 5.**
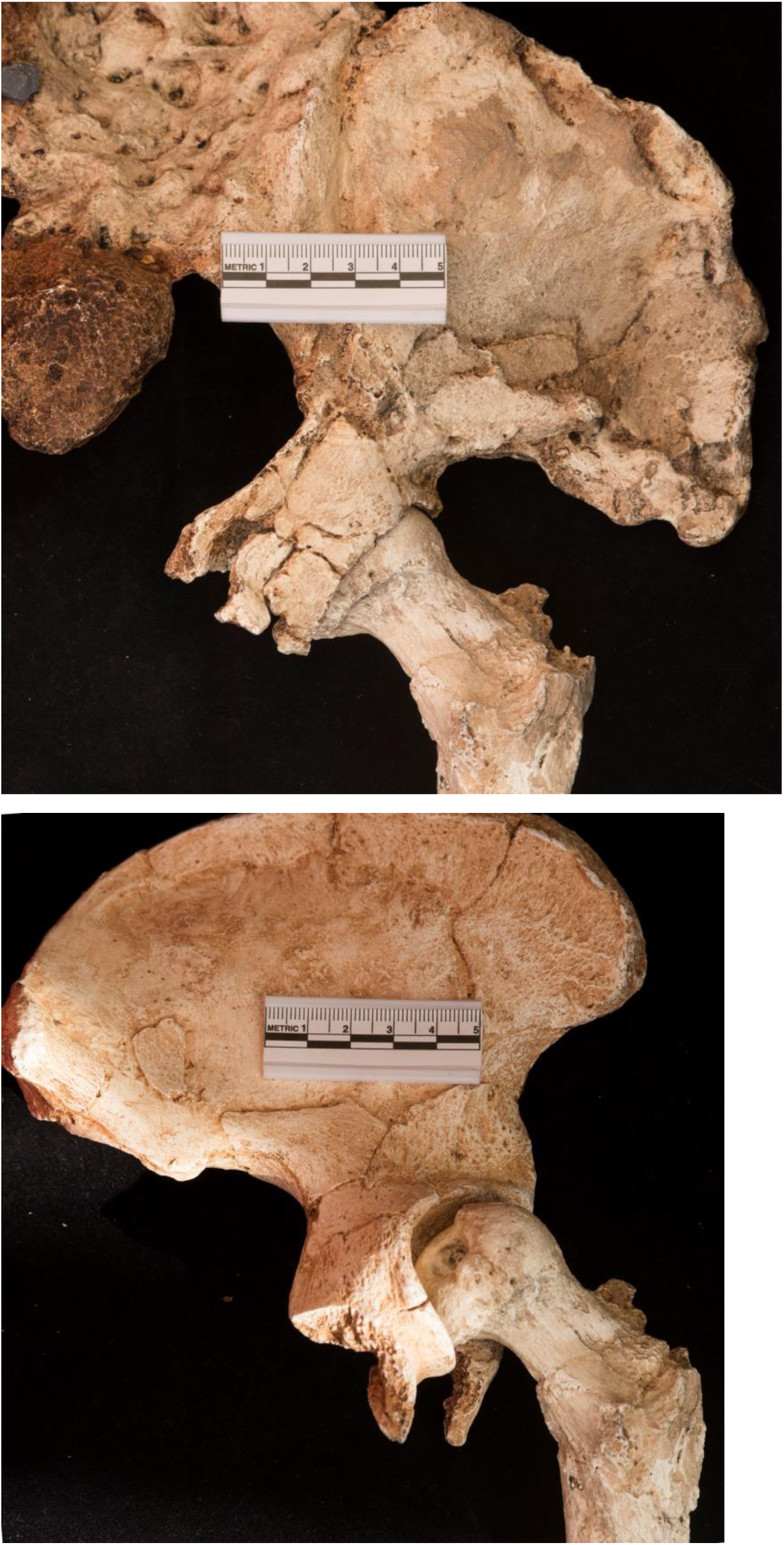
Top, proximal femur of StW 573 mounted in its acetabulum; Bottom, mounted in acetabulum of StW 431 innominate

**Figure 6:**
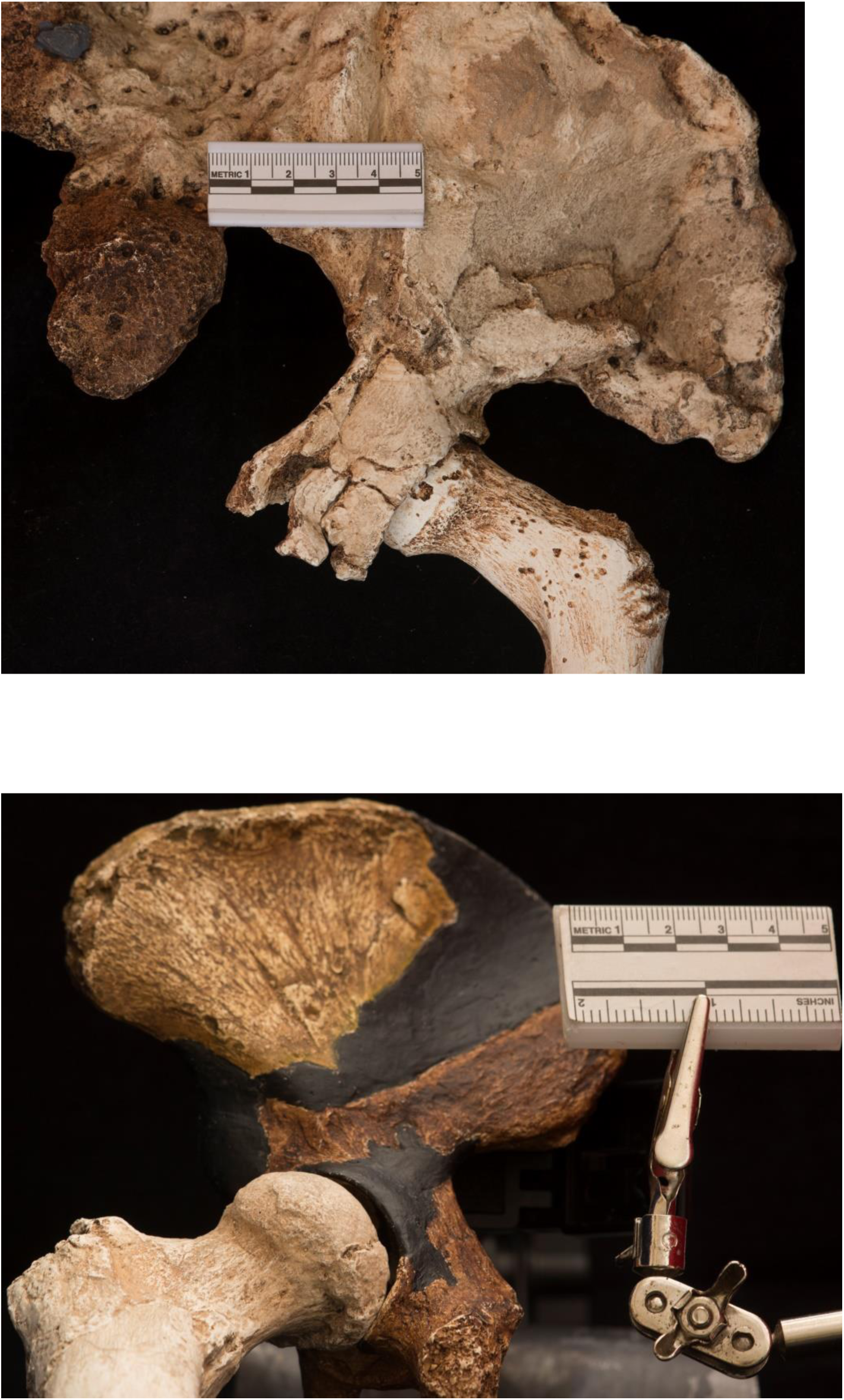
Top, the proximal femur from Jacovec Cavern StW 598 mounted in acetabulum of StW 573; Bottom, the proximal femur of StW 573 mounted in acetabulum of the Sts 14 innominate

**Figure 7:**
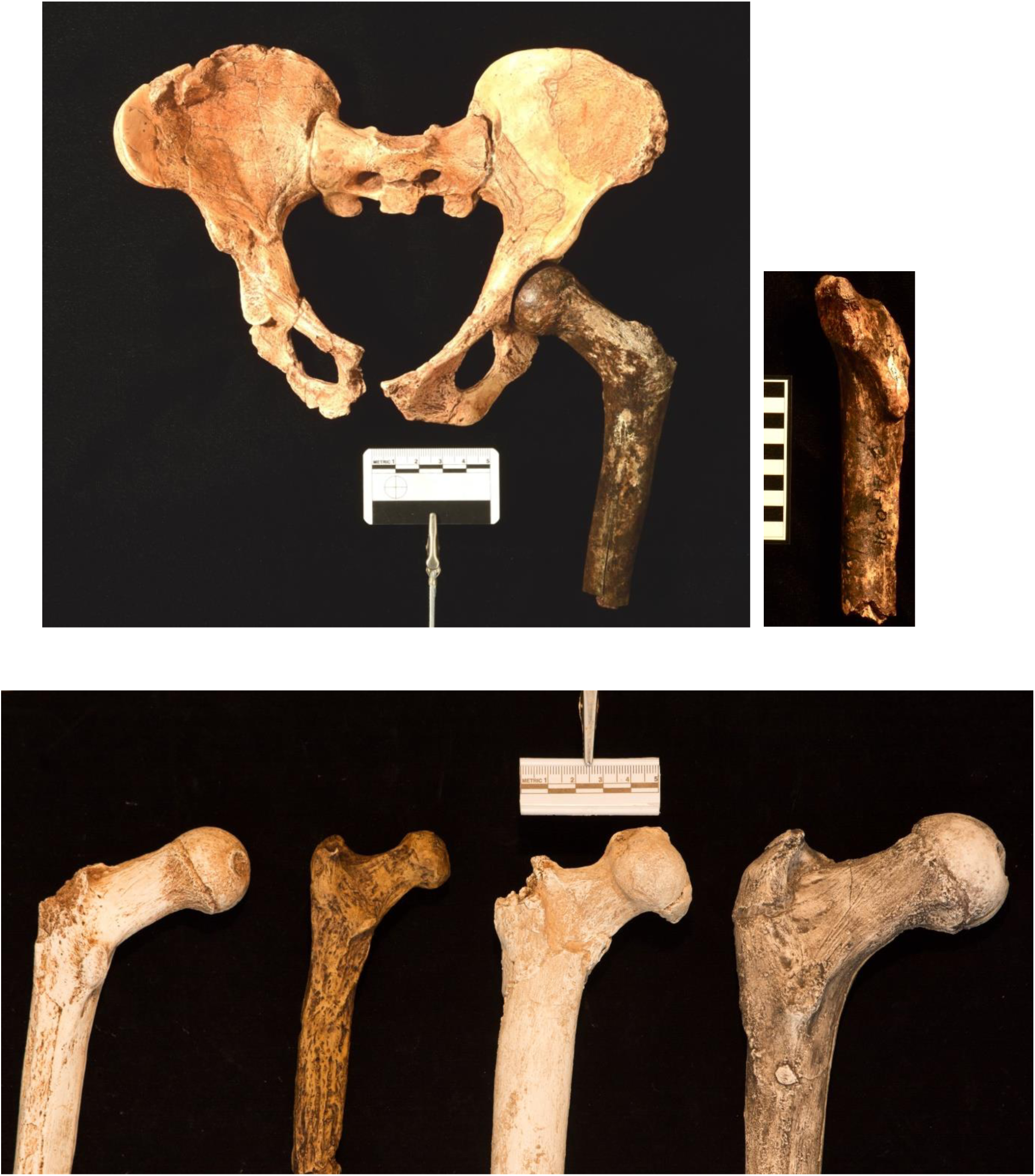
Top left, the proximal femur from Jacovec Cavern StW 598 mounted in the acetabulum of Sts 14; Top right, the StW 367 proximal femur from Member 4; Bottom; Proximal femora of (left to right) Jacovec StW 598, AL-288-1, StW 573 and KNM WT 15000

Fornai et al. (2018) recently reported that StW 431 also differs markedly from Sts 14 in sacral shape, and, like ourselves, they suggest that functional morphs exist in South African *Australopithecus*. In this respect, it is interesting that Toussaint et al. (2003) noted very different body mass estimates for StW 431--42.5 kg using a hominoid RMA regression line, and 41.1 kg using a human regression line, but only 33.4 kg and 22.6 kg respectively on the basis of the lumbosacral region. There are clearly major biomechanical distinctions in the lumbosacral and hip regions, key for the effectiveness of upright walking, between universally recognised *A. africanus* (*eg*., Sts 14) and both StW 573 and StW 431. Not all of them can be put down to a simple relationship to body size – although body size itself is a major difference. Since a palaeospecies is identified by morphological (and one would hope, functional) distance, we argue that the balance of evidence is now strongly in favor of broad recognition of *A. prometheus* as a species distinct from *A. africanus*.

### 2.5 Cranial and Dental Anatomy and Diet

An overall cranial shape similarity is evident with the Bouri Hata hominin, ca. 2.5 Ma, (Asfaw et al. 1999), but some aspects of cranial morphology suggest to Clarke and Kuman (2018, submitted) that an ancestral relationship of StW 573 to *Paranthropus* may be possible. Beaudet et al. (2018a, in press) conclude from the remarkably well preserved endocast that the brain was small (perhaps surprisingly so) and undistinguished from that of other non-human great apes (NHGAs).

Although it will take some time for the mandible to be detached safely from the cranium, microCT scanning has revealed that wear distribution as well as dental arcade shape resemble those of Kanapoi *A. anamensis,* 4.17-4.12 Ma., where Ward *et al.* (2001, p. 351) found that the ‘teeth exhibit a distinctive pattern of wear. Evident in older individuals, the anterior teeth are worn very heavily, much more so than the molars and premolars.’ Ward et al. (2001) cite evidence that *A. anamensis* was taking a tough C4 diet, which might suggest open environments. But faunal analysis suggested to Reed (1997) that Kanapoi paleoenvironments at the time of *A. anamensis* were closed woodland. However, Behrensmeyer and Reed (2013) note that other evidence, including stable isotopes, possibly non-arboreal monkeys and micromammals, and characteristics of paleosols, suggest that open habitats also existed. Similarly, Cerling et al. (2013) found that (δ^13^C) stable isotopes in dental enamel of *A. anamensis* suggest a C3-dominated diet (leaves and fruits from trees and shrubs, etc.). Further, comparative evidence from extant colobines (Koyabu and Endo, 2010) indicates that similar wear distribution may result from consumption of tough-skinned arboreal fruit. Of course consumption of tough-coated arboreal fruit and consumption of tough-coated terrestrial resources (such as corms and tubers) are not mutually exclusive. The Woranso-Mille hominin KSD-VP-1/1, now dated to some 3.6 Ma (Haile-Selassie, 2016) and attributed to *A. afarensis*, also appears from faunal evidence to have occupied a primarily wooded environment (Su, 2016), with browsers dominant and grazers a relatively small component, but with some aquatic species, such as crocodiles and an otter, *Torolutra*, suggesting that the locality samples a riverbank community. δ13C determinations from *A. afarensis* at Woranso-Mille suggest a balance of C3 and C4 items in diet (Wynn et al., 2013; Levin et al., 2015), but microwear (Ungar et al. 2010) closely resembles that in *A. anamensis*.

## 3. Functional Interpretation

As noted above, morphometric and general anatomical descriptions of StW 573 long bones are provided by Heaton et al. (2018, submitted); of the scapula by Carlson et al. (2018, in prep.), of the hand by Jashashvili et al. (2018, in prep.), and the foot by Deloison (2004). Detailed descriptions and morphometrics should be sought therein, as here we restrict our attention to the significance of the postcranial anatomy of StW 573 for locomotor ecology of early hominins. We focus our comparative attention primarily on *A. anamensis* from Kanapoi and KSD-VP-1/1, as they bracket StW 573 in time and are of similar size. StW 431 is younger than StW 573, but we regard this specimen as conspecific. Some comparisons will be made to Sts 14 and AL-288-1, but these specimens are considerably smaller and they are more likely to be adaptively different, quite possibly being (in the case of Sts 14 at least) more arboreal. We do not refer extensively to the considerably later *A. sediba*. As noted by Lovejoy et al. (2016), its forelimb seems curiously derived towards some kind of suspensory locomotion and/or feeding, and its hindlimbs are not reliably reconstructed, the lower limb length having been assumed by Berger (eg. 2013) to equal that of proximal and distal fragments plus the length of the empty matrix between them in situ, despite the likelihood that taphonomic events would affect such an indirect length estimate. We make broad comparisons to the well preserved skeleton of *Homo ergaster*, KNM WT 15000 (Walker and Leakey, 1993), ca 1.5 Mya., as appropriate.

### 3.1. Thorax and Pectoral Girdle

The ribs and vertebral column are currently under study by our team but it appears that the thoracic inlet is narrow, unlike the penecontemporaneous KSD-VP-l/1. This does not support the generalization of Lovejoy *et al*. (2016) from KSD-VP-l/1 that early hominins had abandoned the superiorly narrow ribcage typical of NHGAs. On the other hand, the clavicles (Figure 9 top) are broadly humanlike in form, and indeed remarkably long, very similar to those in the much taller KNM WT 15000 *Homo ergaster*. The right clavicle is complete (Figure 9 top). Like that of KNM WT 15000 (Figure 9 bottom), it is delicate, with a clear S shape very similar to that exhibited by humans. The strong sigmoid curvatures would increase moment arm for potential stabilizers of the shoulder girdle against the humerus, such as the clavicular head of the pectoralis major, the deltoid, and pectoralis minor. The most remarkable feature, however, is the length: 14 cm, in an early human ancestor estimated to be ca. 130 cm in stature (RJC pers. comm. to RHC). This clavicular length equals typical means for adult humans worldwide (Trinkaus et al., 2014) and is in striking contrast with the short clavicle of *A. sediba* MH2 as reported by Schmid et al. (2013).

**Figure 8.**
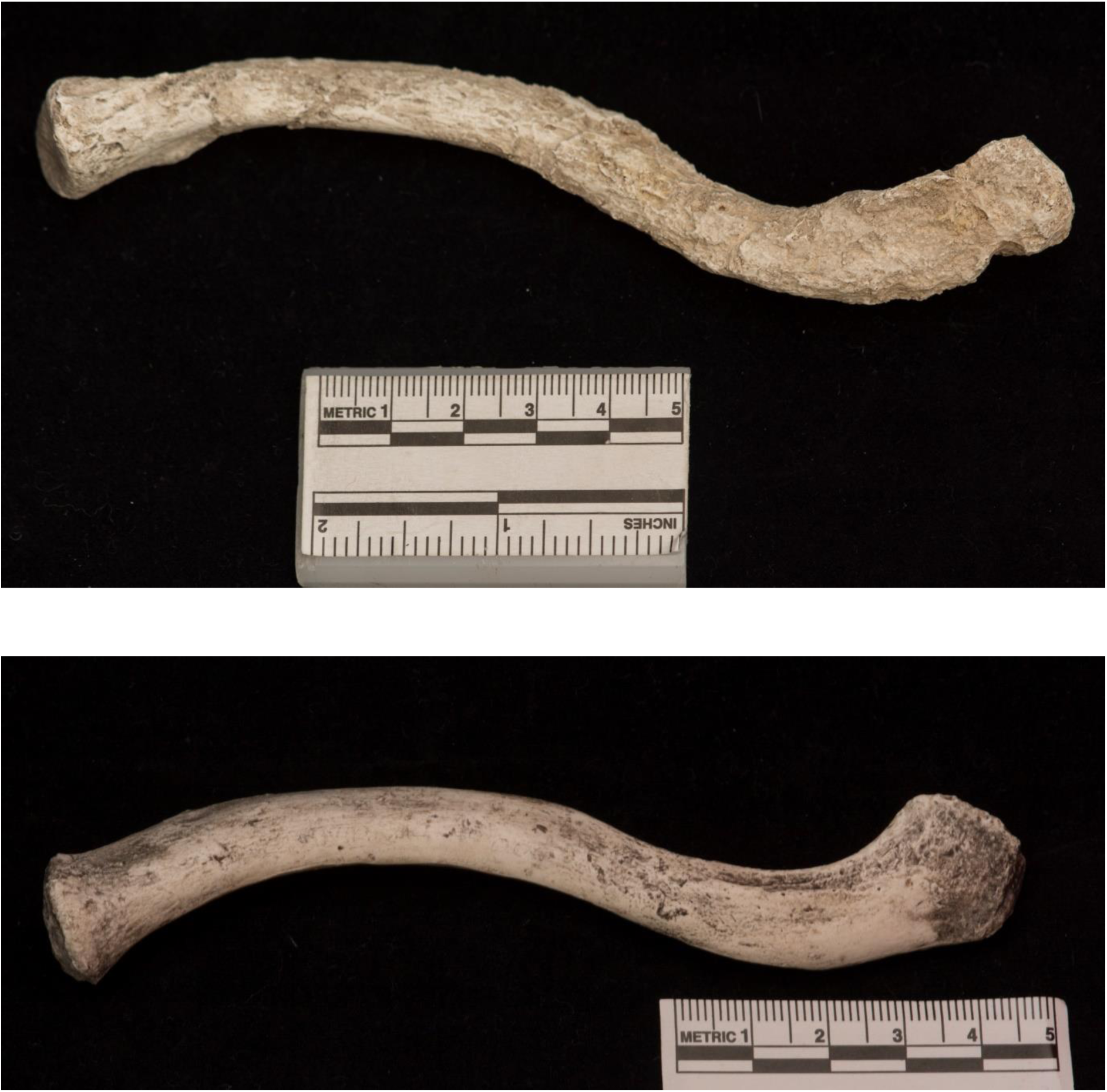
Top: The right clavicle of StW 573 Bottom: The right clavicle of KNM WT 15000

**Figure 9:**
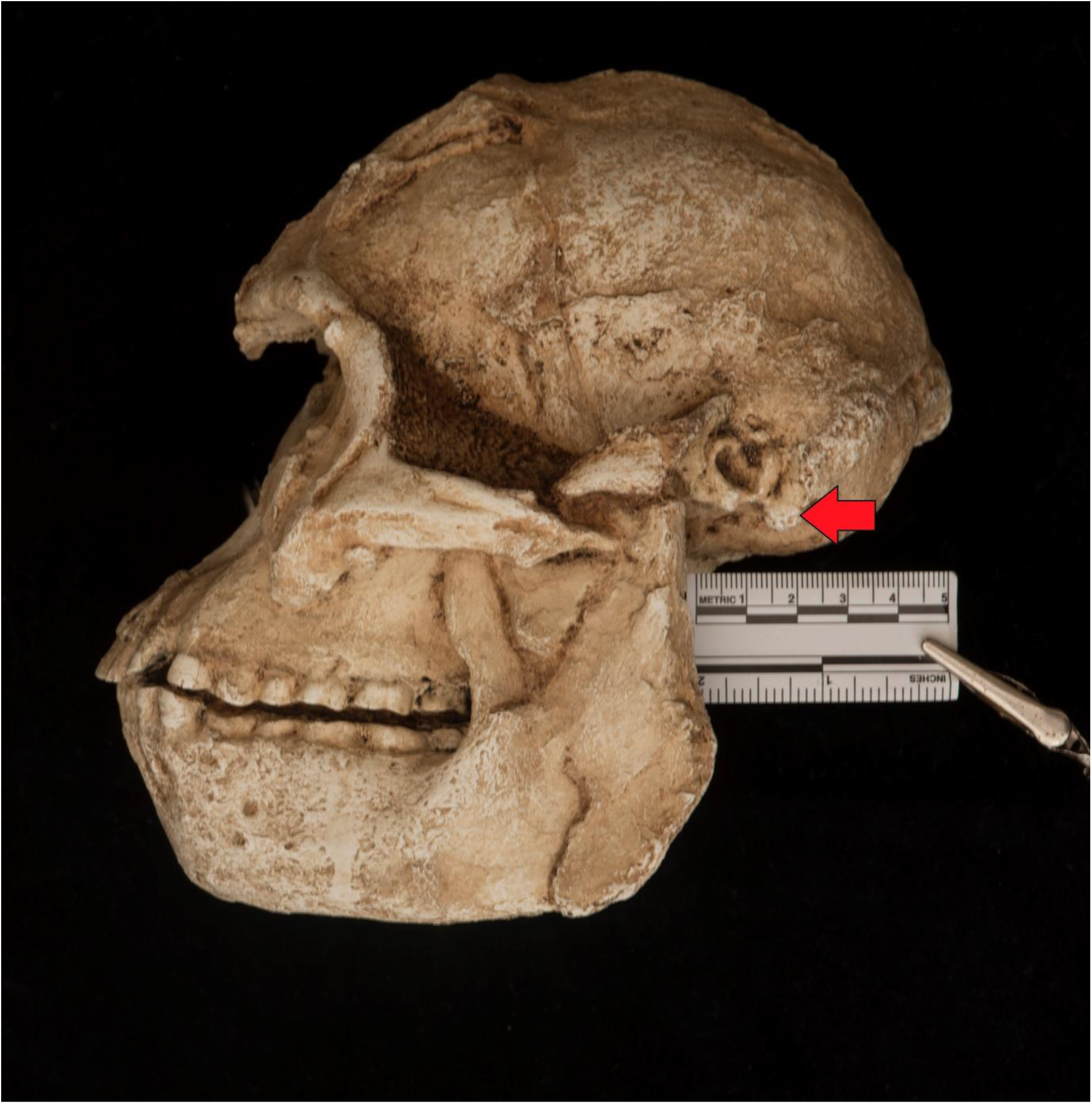
The mastoid process (arrow) on the skull of StW 573

Given the likely close relationship of the male StW 431 to StW 573, we virtually bisected the reconstructed articulated os innominatum and sacrum through the sacral midline and mirrored it. The bi-iliac width of the reconstructed StW 431 pelvis (Fig. 1, bottom) was thus estimated at 30 cm. Given the dimensional similarities between the StW 573a and StW 431 pelves, the bi-iliac breadth of StW 573 cannot have been much less than the 30 cm. biiliac distance in StW 431, which compares to mean values in modern human females of around 28 cm. (see eg. Simpson et al., 2008). Since StW 573’s clavicle was 14 cm long, and assuming some 3 cm inter-clavicular distance (we lack a sternum), her bi-acromial distance would have been some 28-30 cm, very similar to the likely bi-iliac breadth, suggesting that the trunk was more or less of equal width superiorly and inferiorly, unlike the ribcage. This mismatch between a narrow thoracic inlet and broad shoulders suggests the latter was the subject of active selection for large moments at the glenohumeral joint, and hence powerful climbing. A preliminary canonical variates plot of scapular geometry by Carlson et al. (2018, in prep.) based on MicroCT and virtual reconstruction of the shattered scapular blade shows that StW 573 occupies a position very close to MH2 *A. sediba,* but also close to KSD-VP-1/1, *Gorilla* and *Pongo.* However, plots for *Pan* and *Homo* (particularly KNM-WT 15000) lie quite distant from StW 573, at the left and right extremes of the plot. The glenoid fossa is certainly more cranially oriented than in *Homo.* Either way, the geometry of the pectoral girdle of *A. prometheus* does not seem to resemble the ‘shrugged’ girdle proposed by Churchill et al. (2013) for *A. sediba.* Weak expression of the mastoid process on the skull of StW 573 (Figure 10) indicates that the sternocleidomastoid was by no means as powerful as would be expected with such a ‘shrugged’ posture. Indeed, the distinction between the short clavicle of *A. sediba* (1.97 Ma) and the long clavicle of the much earlier *A. prometheus* (3.67 Ma) suggests that any elevated pectoral girdle posture in *A. sediba* is derived, not ancestral as claimed by Churchill et al. (2013). Following Rein et al. (2017), we must consider whether suspensory performance was selected for in *A. sediba*, possibly in connection with postural feeding adaptations.

**Figure 10:**
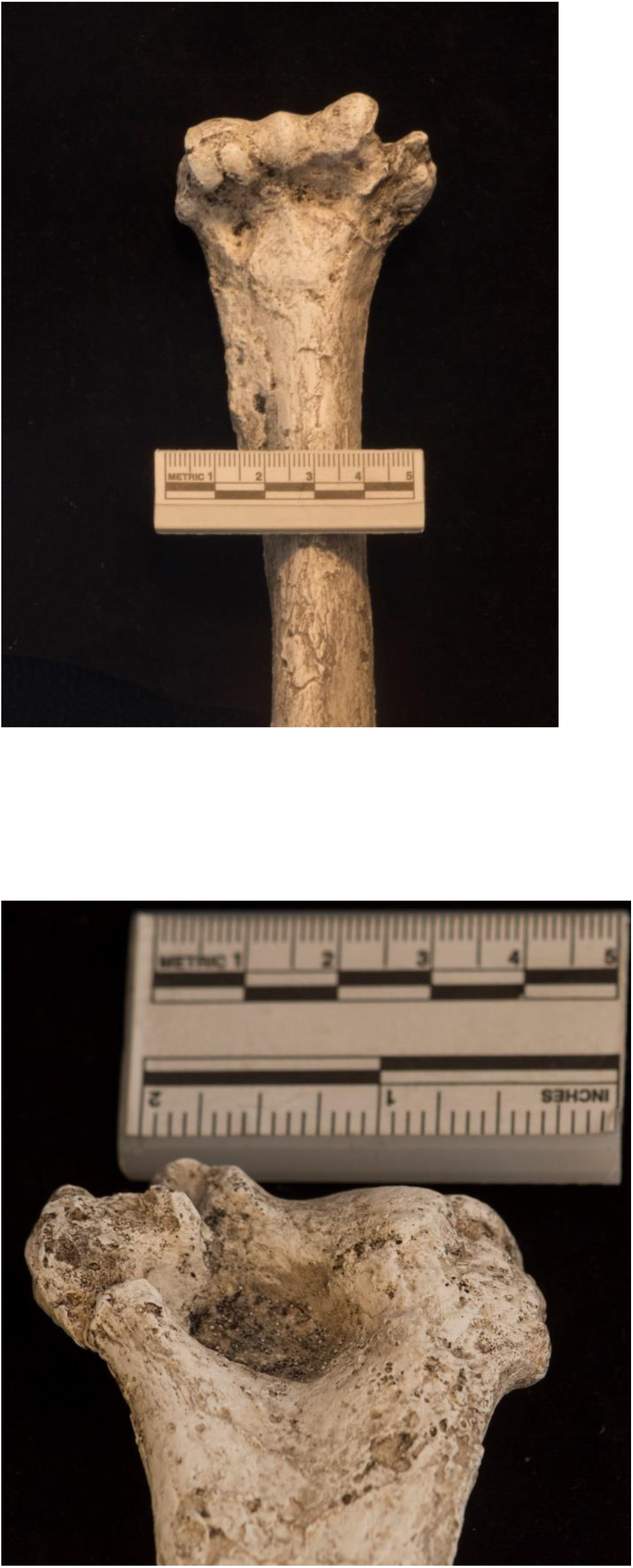
Distal humerus of StW 573. Top, ventral view showing brachioradialis crest; Bottom, dorsal view showing shape of distal condyles

### 3.2 Arm

The right humerus is crushed but intact. It is articulated proximally with the scapula and distally with the radius and ulna. The head of the detached left humerus is crushed and so the size of the deltoid tuberosity cannot be assessed. Muscle markings are moderately strong, particularly the intact brachioradialis crest (Figure 10 top), which appears substantially larger than the damaged crest in KSD-VP-1/1b figured by Lovejoy et al. (2016). This implies more power in pronation in StW 573 (which hypothesis again can be tested in silico). The distal humeral condyles (Figure 10 bottom) appear very similar in form to those of the Kanapoi *A. anamensis* KNM-KP 271, figured by Hill and Ward (2018) in having, for example, a more salient lateral margin for the trochlear articulation than KNM WT 15000 (Figure 11). This feature might imply less axial ‘rocking’ of the ulna than occurs in our genus, but as Lovejoy and colleagues (2016) note, these distinctions are not so major as to necessarily imply active selection. And, as Hill and Ward (1988) note, distal humeral morphology is very variable in humans. Further, Hill and Ward (1988) comment that the Kanapoi distal humerus shows a clear fracture and needs to be considered with caution.

**Figure 11:**
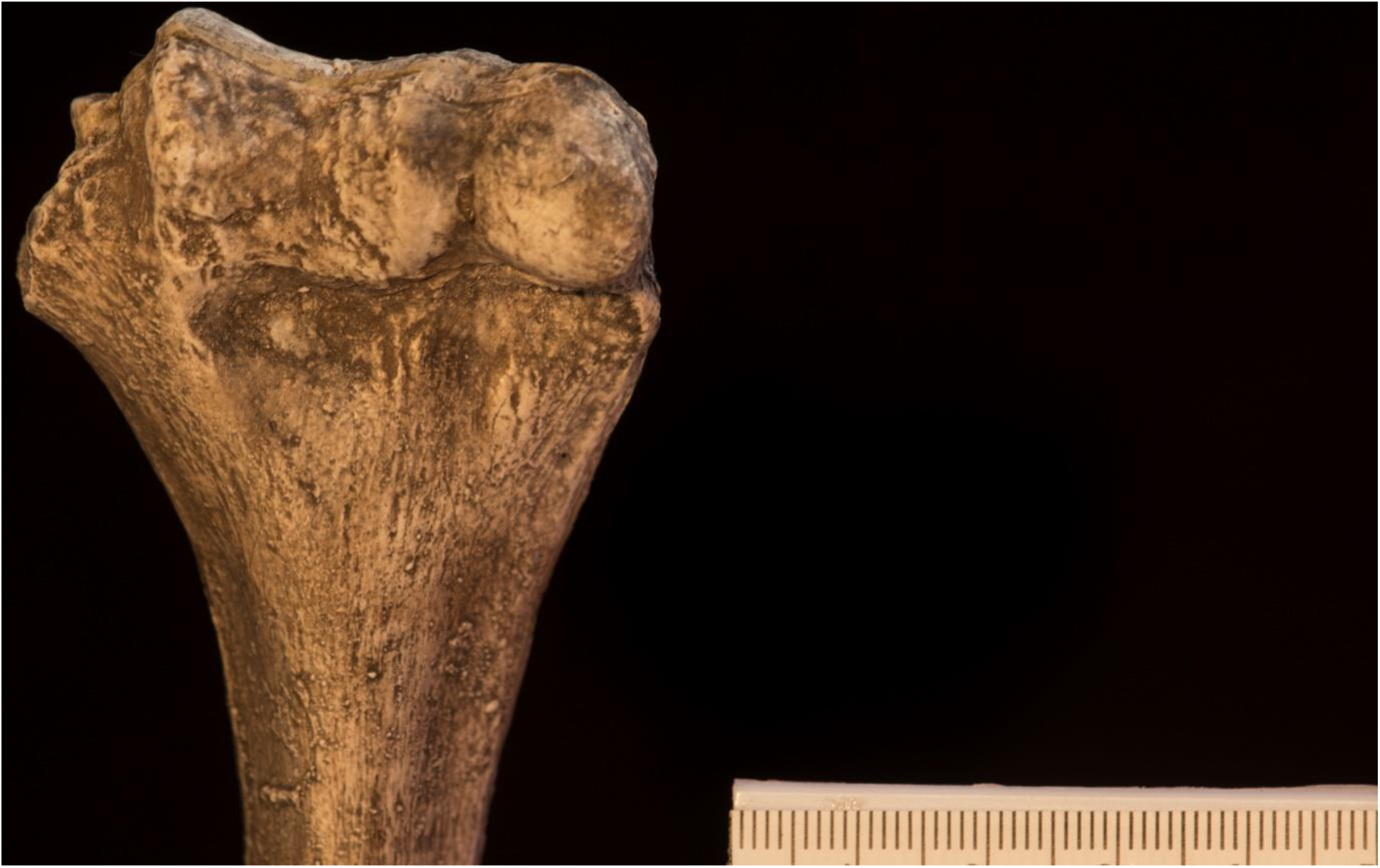
Distal humerus of KNM WT 15000, dorsal view

With regard to the ulna, the shape of the StW 573 trochlear notch agrees more closely with the human ulna figured in Lovejoy *et al.* (2016) than with either KSD-VP-1/1 or AL-288-1 in its somewhat less anterior orientation. Following those authors, we refrain from functional interpretation at this moment. Shaft curvature appears more marked than that figured by Lovejoy et al. (2016) for KSD-VP-1/1, but it appears from Drapeau et al. (2005) that curvature is variable in early hominins. There is no radius for KSD-VP-1/1, but that of the Kanapoi *A. anamensis*, as figured by Ward et al. (2001), is both similar in morphology and near-identical in length to that of StW 573. Retention of such a long radius (see section 3.5, *Limb proportions*), especially in combination with a relatively powerful brachioradialis, implies power in flexed/pronated elbow postures, most likely employed during climbing.

### 3.3. Pelvic girdle

The os innominatum of both StW 573 and StW 431 corresponds broadly with the form shown by Lovejoy et al. (2016, Fig 8.21) for *A. afarensis*, with both a greater sciatic notch and anterior inferior iliac spine evident, although the latter has sheared off in StW 573. We need not refer further to the pelvic girdle until the crushed pelvis of StW 573 has been restored by retrodeformation. Hence most information is drawn from StW 431 (see Toussaint *et al*., 2002 and Kibii and Clarke, 2003). However, we should note that as Kozma et al. (2018, p. 1) pithily conclude from a study of hip extensor mechanicsm, ‘*Ardipithecus* was capable of nearly human-like hip extension during bipedal walking, but retained the capacity for powerful, ape-like hip extension during vertical climbing. Hip extension capability was essentially human-like in *Australopithecus afarensis* and *Australopithecus africanus,* suggesting an economical walking gait but reduced mechanical advantage for powered hip extension during climbing.’ Contra Lovejoy et al. (2016) who unequivocally attribute a short ischium in *Homo* to running, Kozma et al. (2018) demonstrate that a short ischium greatly enhances distance travelled for energy consumed in walking. But it is worth noting that musculoskeletal modelling by some of us (Goh et al., 2017) showed that in terms of joint moments and torques exerted by all major lower limb extrinsic muscles, the ability of gorillas to walk bipedally is not limited by their adaptations for quadrupedalism and vertical climbing.

### 3.4 Femur, Tibia, Hip, Knee and Ankle

We have noted that the femoral head of StW 573 is large, and the femoral neck is short compared to *A. africanus* sensu stricto (e.g., the proximal femur from Jacovec Cavern StW 598 [Partridge et al., 2003]) and *A. afarensis* AL-288-1. In that respect it resembles KNM-WT 15000 more closely. Unfortunately, there is as yet no proximal femur for the more size and age-matched *A. afarensis* KSD-VP-1/1. The left distal femur of *A. afarensis* KSD-VP-1/1 is poorly preserved, especially the medial condyle, but Lovejoy et al. (2016) report that the (restored) lateral condyle is ‘elliptical’, and like StW 573, the patellar groove is deep and shows a high lateral wall for patellar retention, as noted by Heaton et al. (2018, submitted).

However, the lateral femoral condyle of StW 573 (Figure 12) is not only posteriorly ‘elliptical’ (to use Lovejoy and colleagues’ [2016] term), but more specifically like humans, has a relatively rounded posterior/dorsal section and flat anterior/ventral section. Again like humans, the medial femoral condyle is more evenly rounded dorsoventrally (Figure 13). The knee of KSD-VP-1/1 does, as Lovejoy et al. (2016) state, appear to show a valgus angle (see their Figures 8.6 and 8.6). But it is, like that of StW 573 (Figure 14) more weakly marked than in KNM-WT 15000 and particularly than in AL-288-1 (Stern and Susman 1983), where, taken at face value, the angle probably reaches an extreme among hominins.

**Figure 12:**
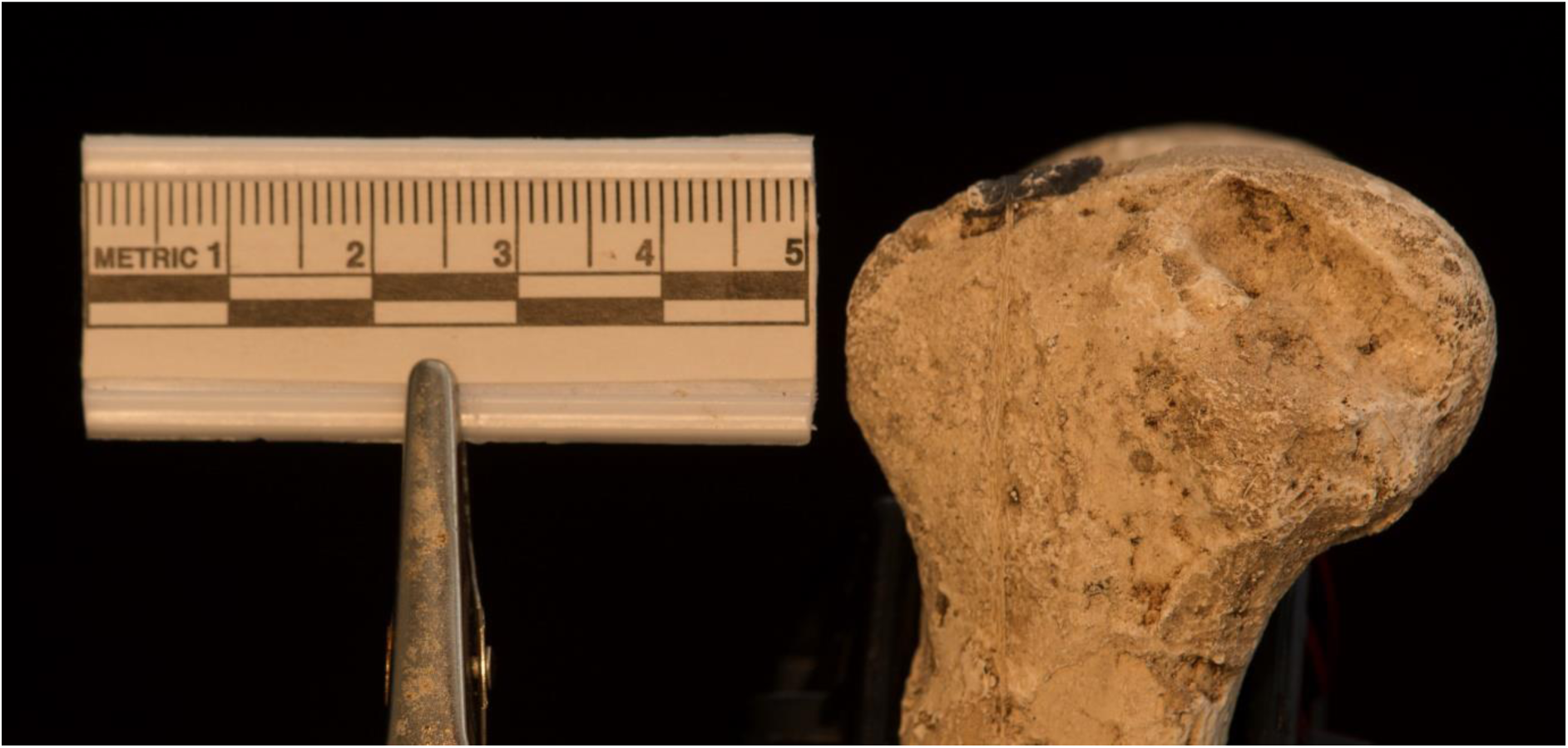
Lateral distal femoral condyle of StW 573

**Figure 13:**
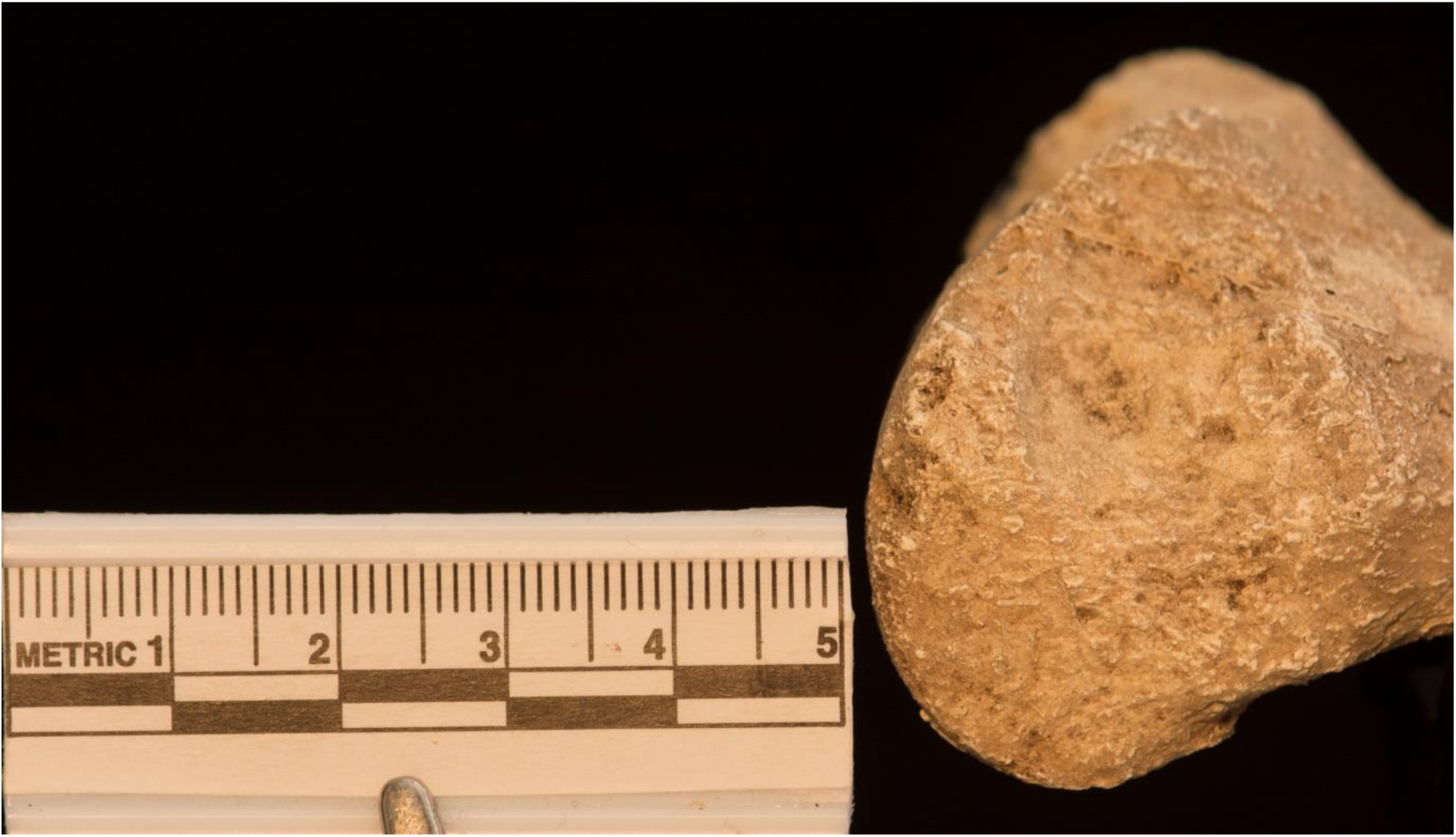
Medial distal femoral condyle of StW 573

**Figure 14:**
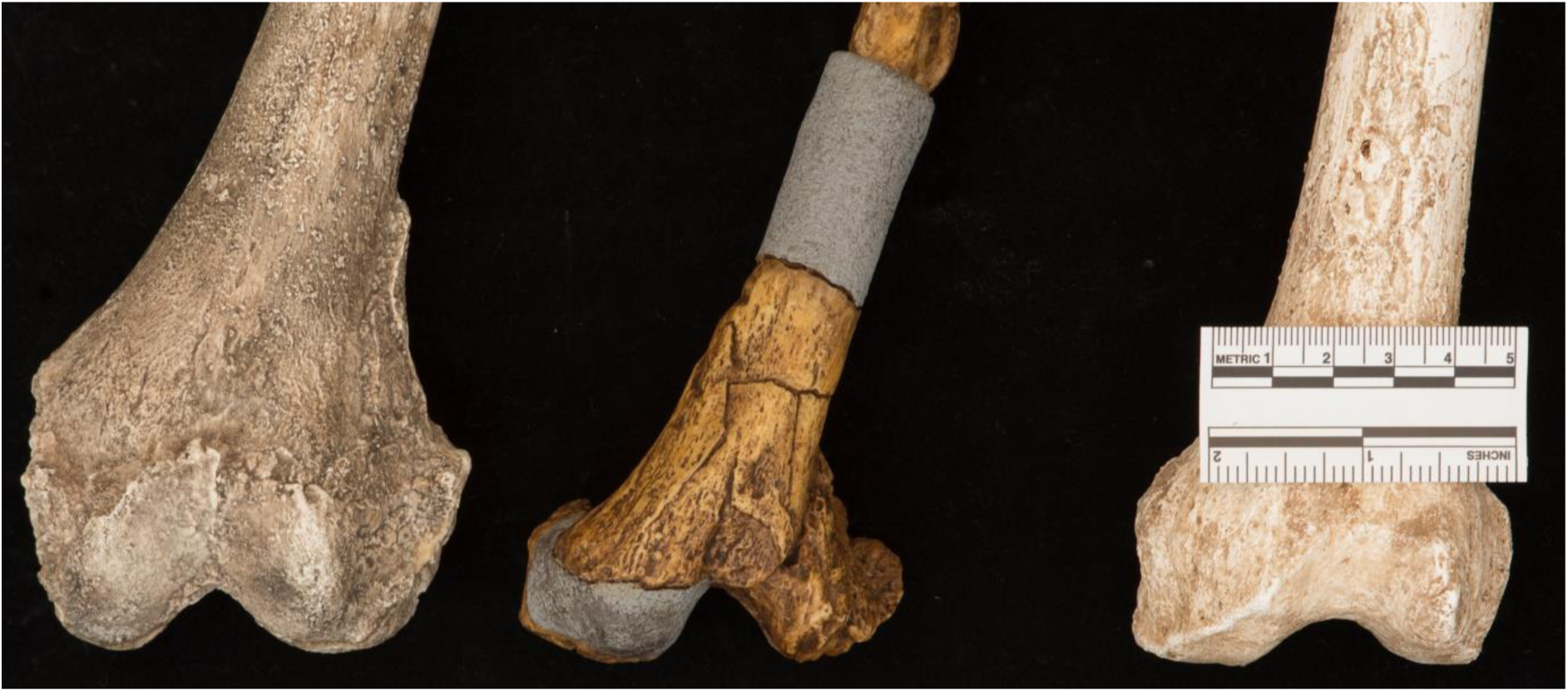
Valgus angles of the knee in, left to right: KNM WT 15000, AL-288-1 and StW 573

There is detailed evidence of the morphology of the proximal surface of the *A. anamensis* tibia from Kanapoi KNM-KP 29285A, 4.16 Ma (reviewed in Ward et al., 2001), which is shown in Figure 15 as a visualization of an stl file (open source, from: africanfossils.org. XYZ dimensions 68.00; 103.30; 60.66 mm.). Ligamentous and muscular attachments are detailed, but although preservation of StW 573 and KNM WT 15000 is excellent, these are not identifiable with any confidence in either specimen (Figure 16 top and bottom). The tibia of KSD-VP-1/1 is heavily damaged throughout and the proximal surface carries little information. From Figures 17, 18 and 19, it is clear that KNM-KP 29285A, StW 573 and KNM WT 15000 all have long, concave condyles on the medial side, and short, less concave condyles on the lateral side, which in KNM-WT 15000 and StW 573 are matched by a long rounded section on the medial femoral condyle but an anteriorly flatter lateral condyle. This is the bony basis of the ‘locking’ or ‘screw-home’mechanism of the knee (see eg. Dye, 1987 and Lovejoy, 2007). The condyles and cruciate ligaments form a four-bar linkage. In knee extension, because of the flatter condylar morphology of the ventral part of the lateral condyles, they cease sagittal rotation motion before the medial condyle, and rollback occurs, compressing the lateral meniscus and further immobilizing the lateral condyle so that a passive coronal rotation results, spiralizing fibres in the cruciate ligaments and stabilizing the knee. This allows standing with minimal expenditure of muscular energy for balance and signifies that early hominins from 4.16 Ma onwards (including both *A. anamensis* and *A. prometheus*) were able to stand upright with enhanced efficiency. ARA VP1/701 *Ardipithecus ramidus* lacks most of the femur, and curiously, the nearly complete tibia is largely unreported (see eg. White et al., 2009), so we cannot assess whether *Ar. ramidus* had this important mechanism, despite Lovejoy’s reference to his own (2007) paper discussing the so-called ‘screw-home’ mechanism reviewed above. But as might be expected the associated distal femoral condyle asymmetry is evident in the morphology of the AL-288-1 distal femur, 3.4 Ma (Figure 20) (and see Stern and Susman, 1983 and Lovejoy, 2007).

**Figure 15:**
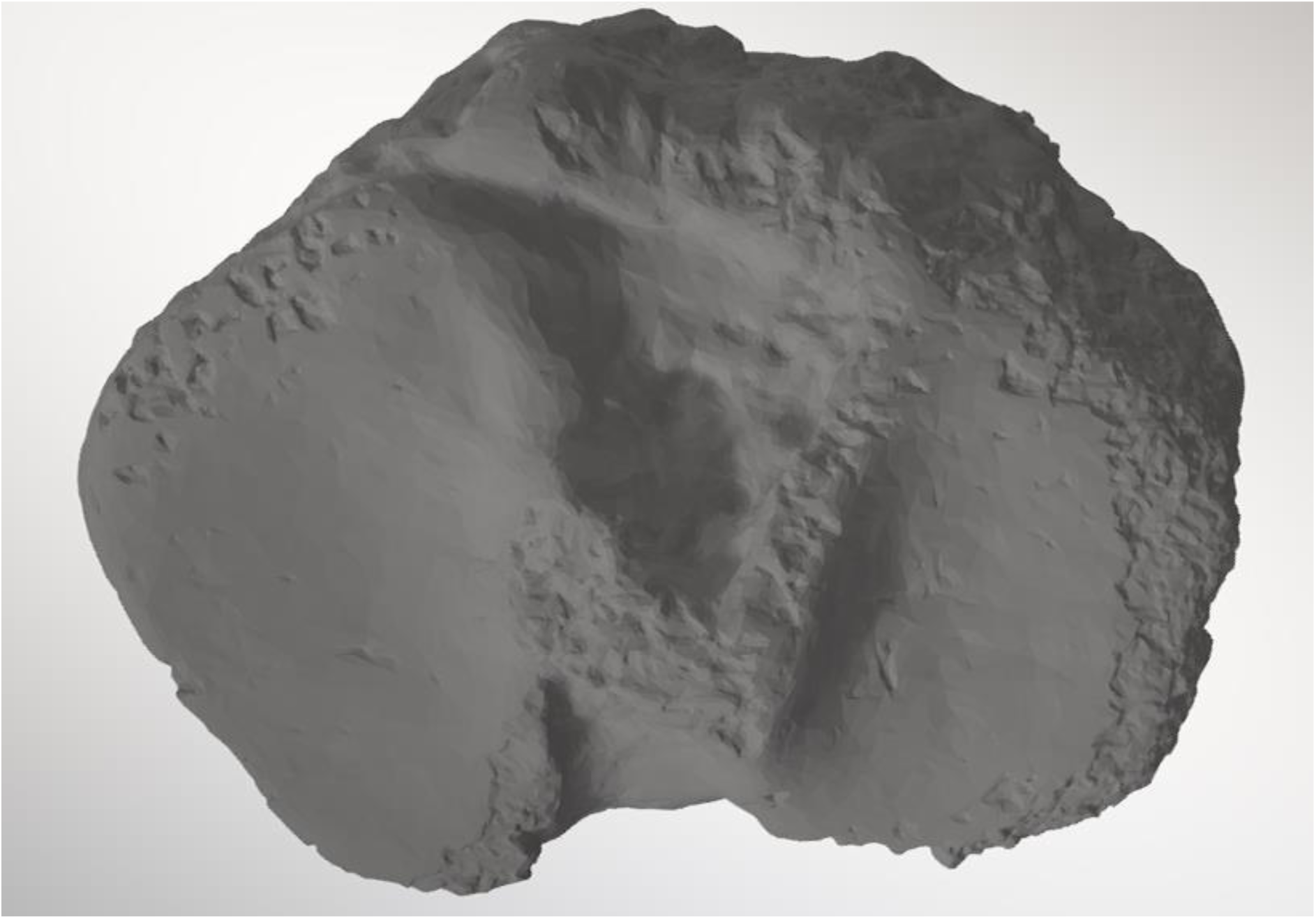
Proximal tibial surface of Kanapoi KNM-KP 29285A (downloaded from open source: www.africanfossils.org, XYZ dimensions 68.00; 103.30; 60.66 mm)

**Figure 16:**
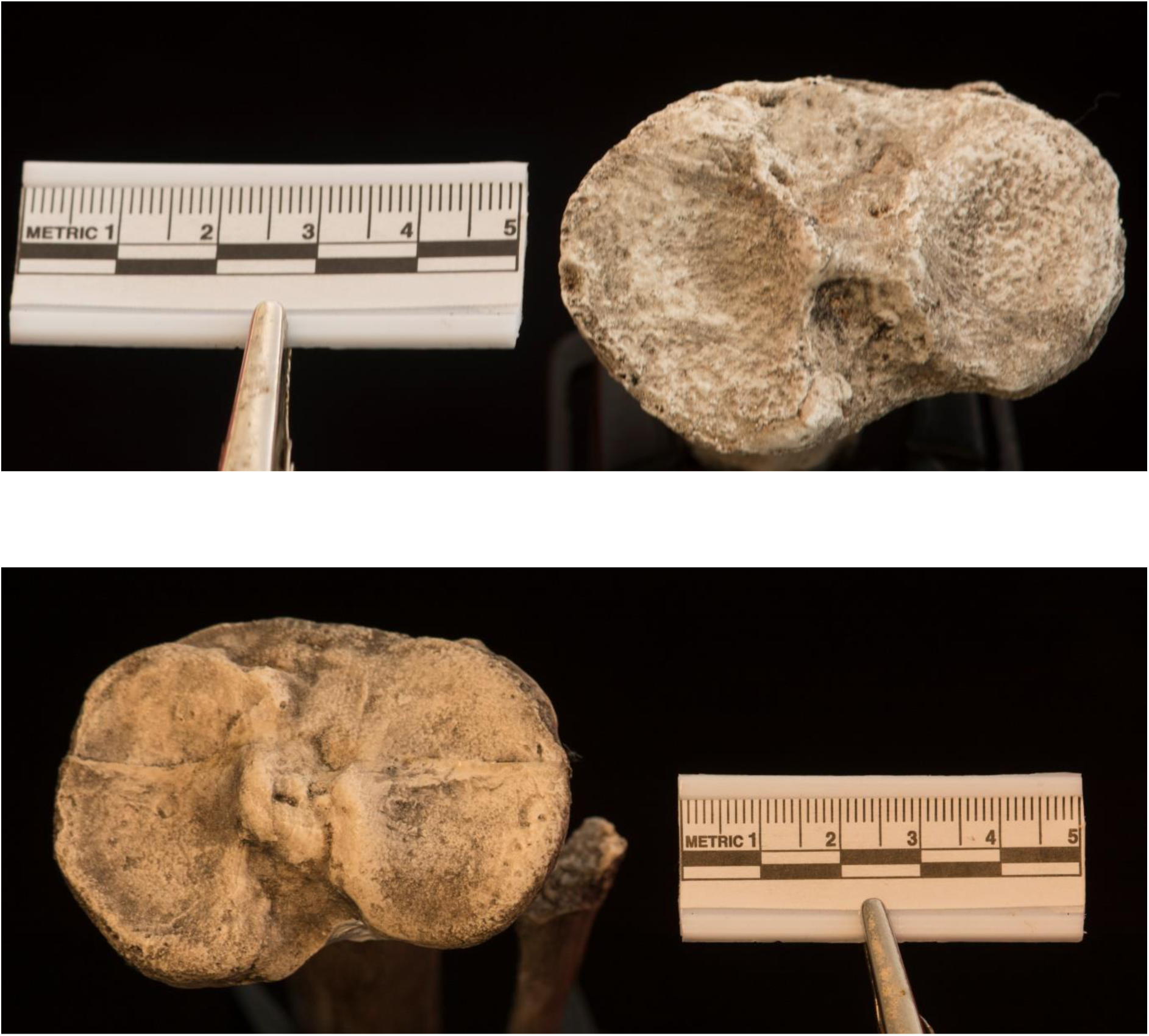
Proximal tibial surface of (Top) StW 573 and (Bottom) KNM WT 15000

**Figure 17.**
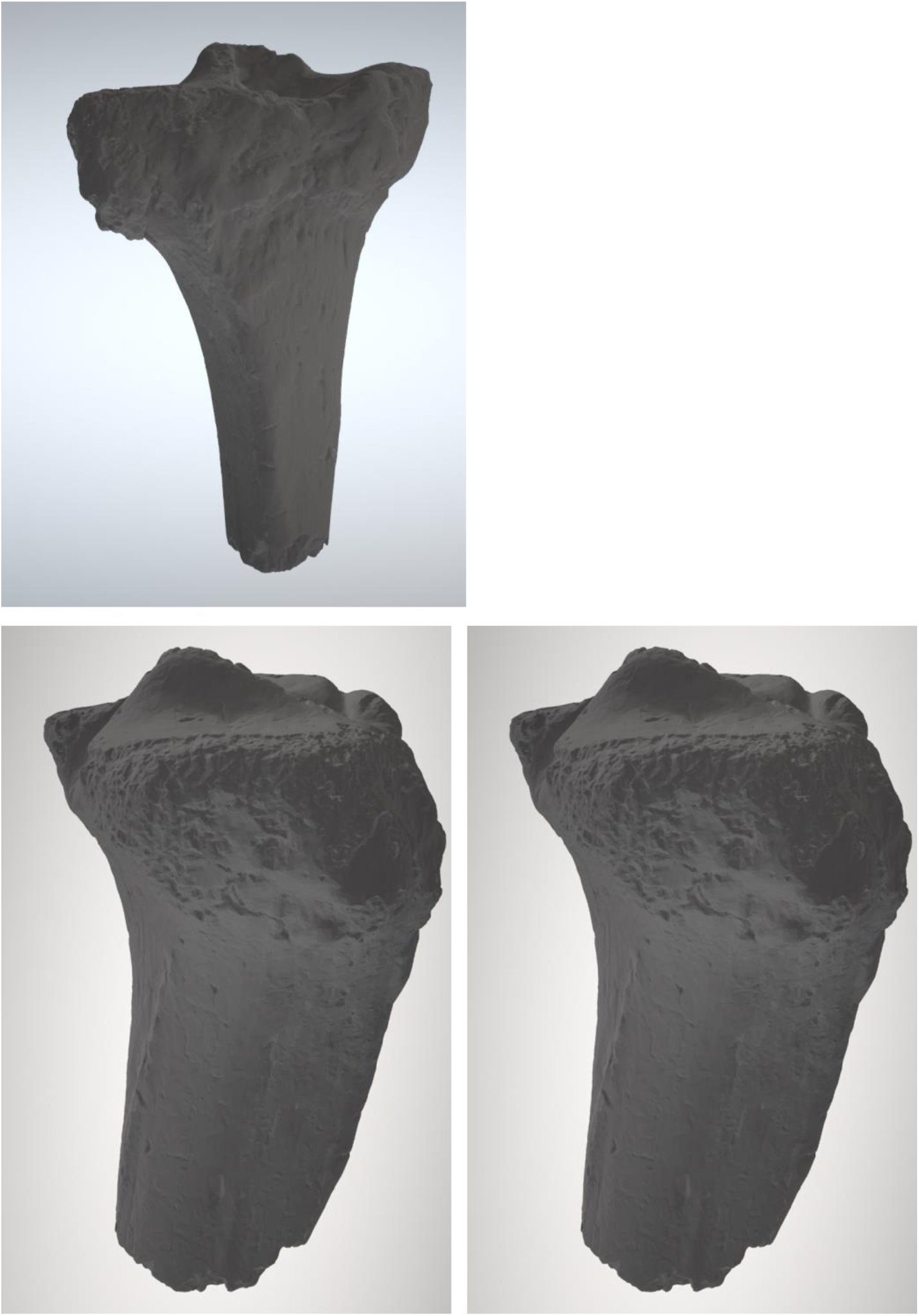
(Top) Frontal, (Bottom left) Lateral, and (Bottom) Medial perspectives of Kanapoi KNM-KP 29285A (downloaded from open source: www.africanfossils.org, XYZ dimensions 68.00; 103.30; 60.66 mm)

**Figure 18.**
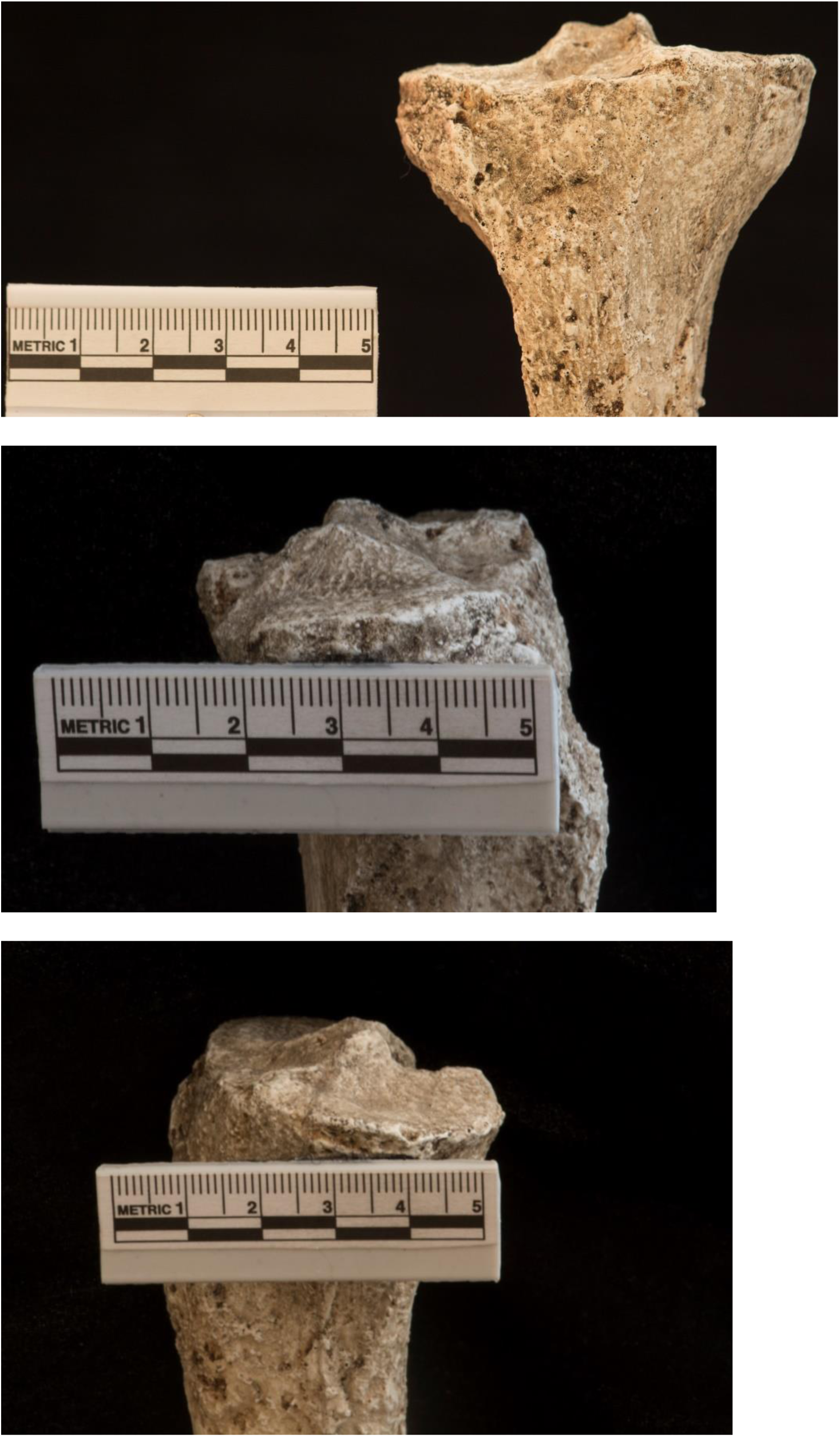
(Top) Frontal, (Middle) Lateral and (Bottom) Medial perspectives of proximal tibial condyles of StW 573

**Figure 19.**
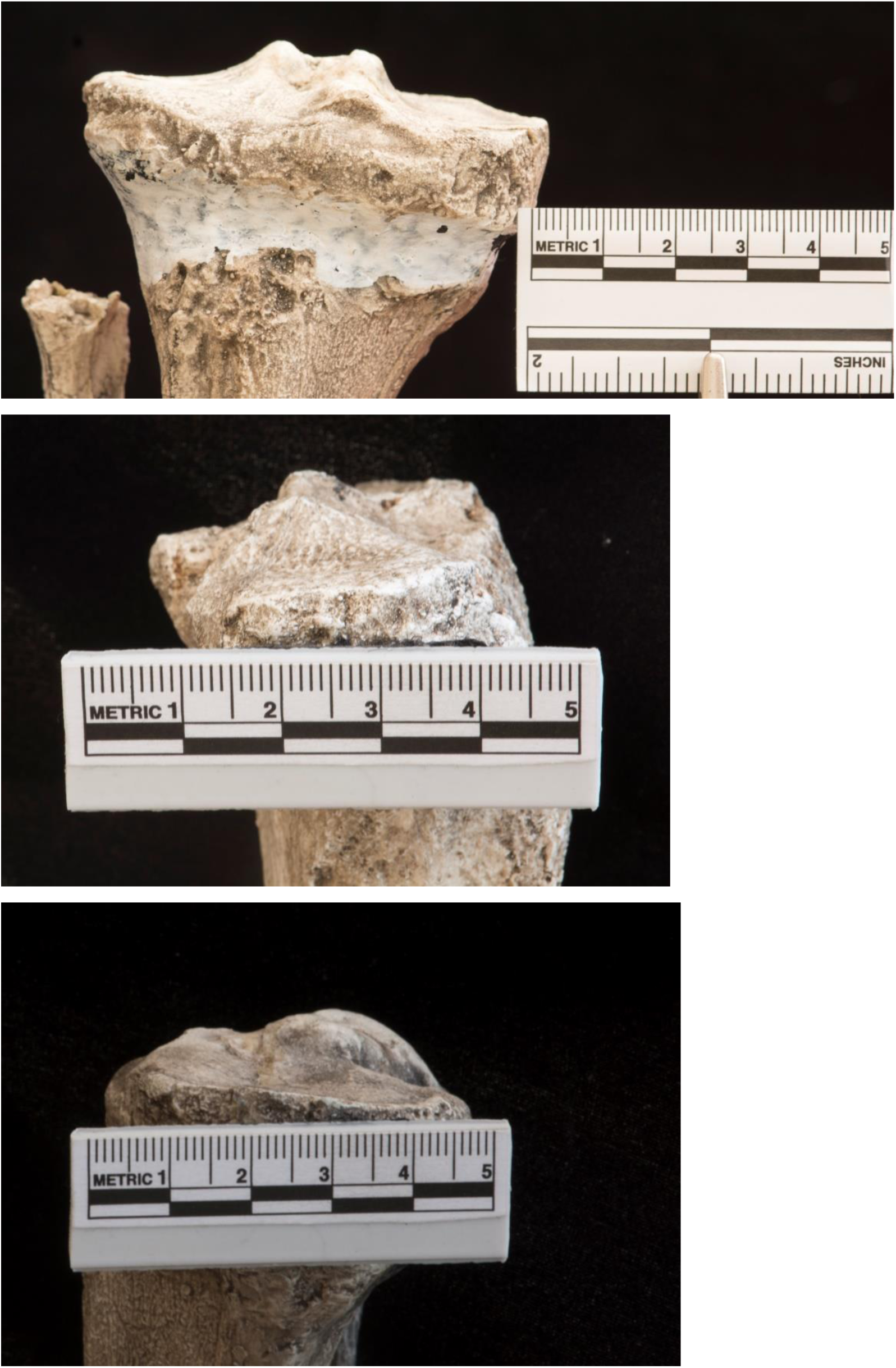
(Top) Frontal, (Middle) Lateral and (Bottom) Medial perspectives of proximal tibial condyles of KNM WT 15000

**Figure 20.**
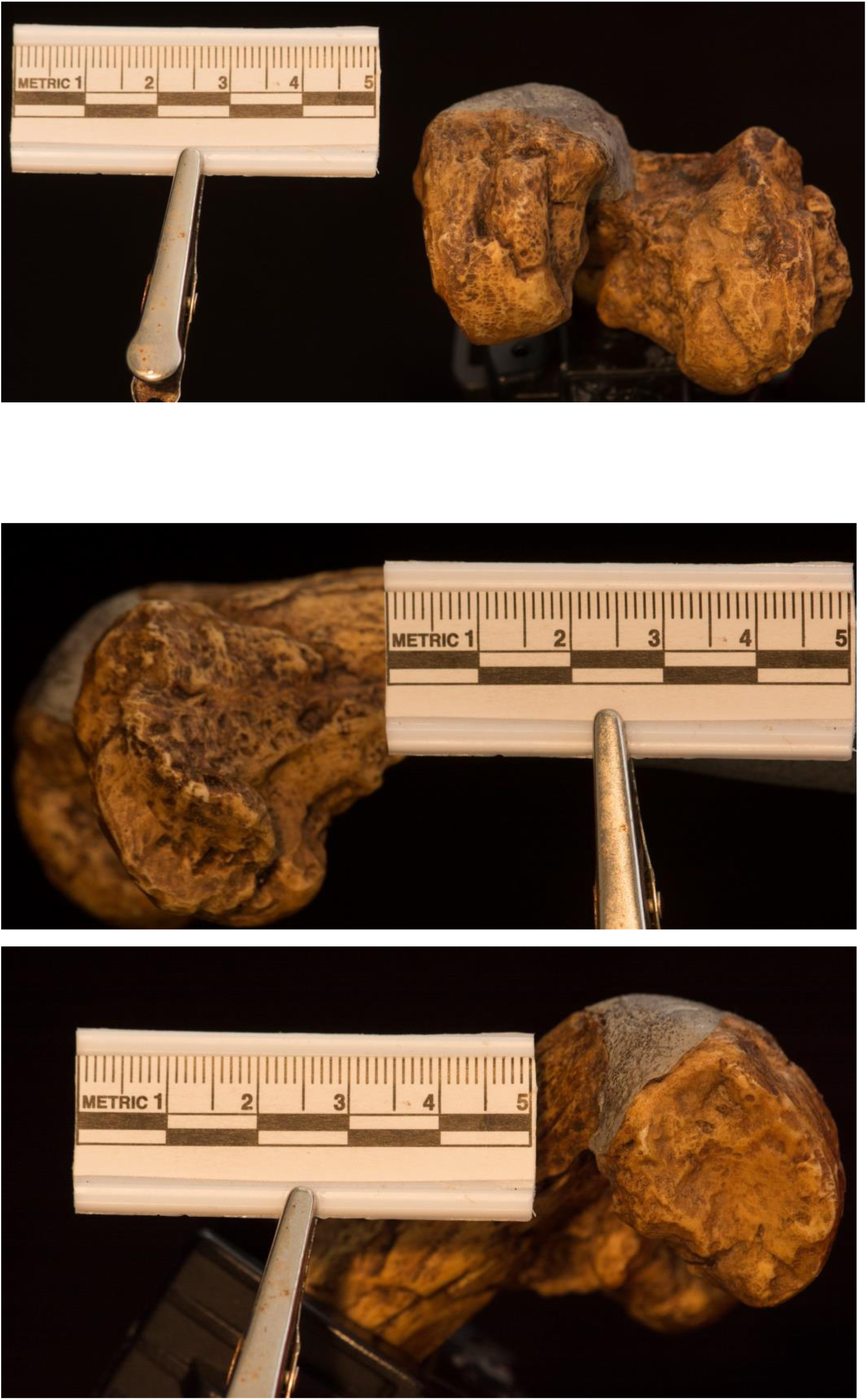
(Top) Axial, (Middle) Lateral and (Bottom) Medial perspectives of distal femoral condyles of AL-288-1

In upright arboreal bipedalism, some of us have shown experimentally (Johannsen et al., 2017) that ‘light touch’ with the fingers on supports between shoulder and waist height significantly enhances balance on unstable supports and reduces thigh muscle activity required to counteract perturbation by some 30%. *Ar. ramidus* could have used this mechanism in upright bipedal walking in the trees, and thus it could have been an effective upright arboreal biped. The same applies to *Pierolapithecus* (12.5-13 Ma, Moyà-Solà et al., 2004).

Lovejoy et al. (2016) draw attention to the short radius of curvature in the talar joint surface of the distal tibia KSD-VP-1/1,, versus the flatter talar joint surface in NHGAs (DeSilva, 2009). Figure 21 shows that the radius of curvature is as short in StW 573 as it is in KNM-WT 15000, and a similarly short radius of curvature can be seen in Figure 22 (an stl model of Kanapoi distal right tibia KNMKP 29285, downloaded from [open source] www.africanfossils.org, Dimensions: x=40.16; y=97.82; z=40.50 mm). Ward *et al.* (2001) note that the maximum concavity of that Kanapoi talar joint surface/plafond is 5 mm. In StW 573 it is ca 4.5 mm, and in KNM-WT 15000 (depending on side) it is also ca 4.5 mm. In each case the shape of the talar joint surface is square, rather than rectangular as tends to be the case in NHGAs.

**Figure 21.**
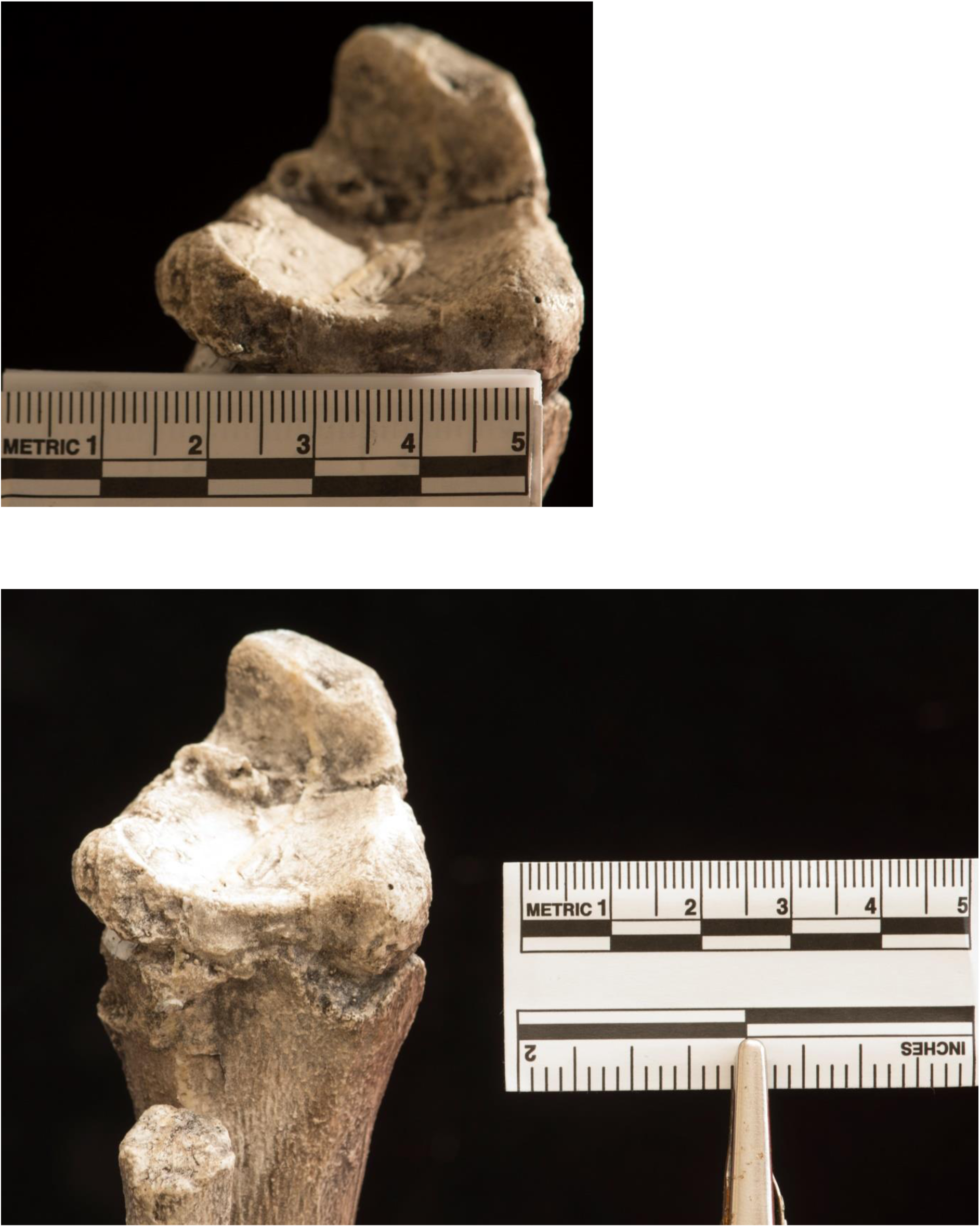
Radius of curvature of the distal tibial condyle/plafond Top: StW 573 Bottom: KNM WT 15000

**Figure 22.**
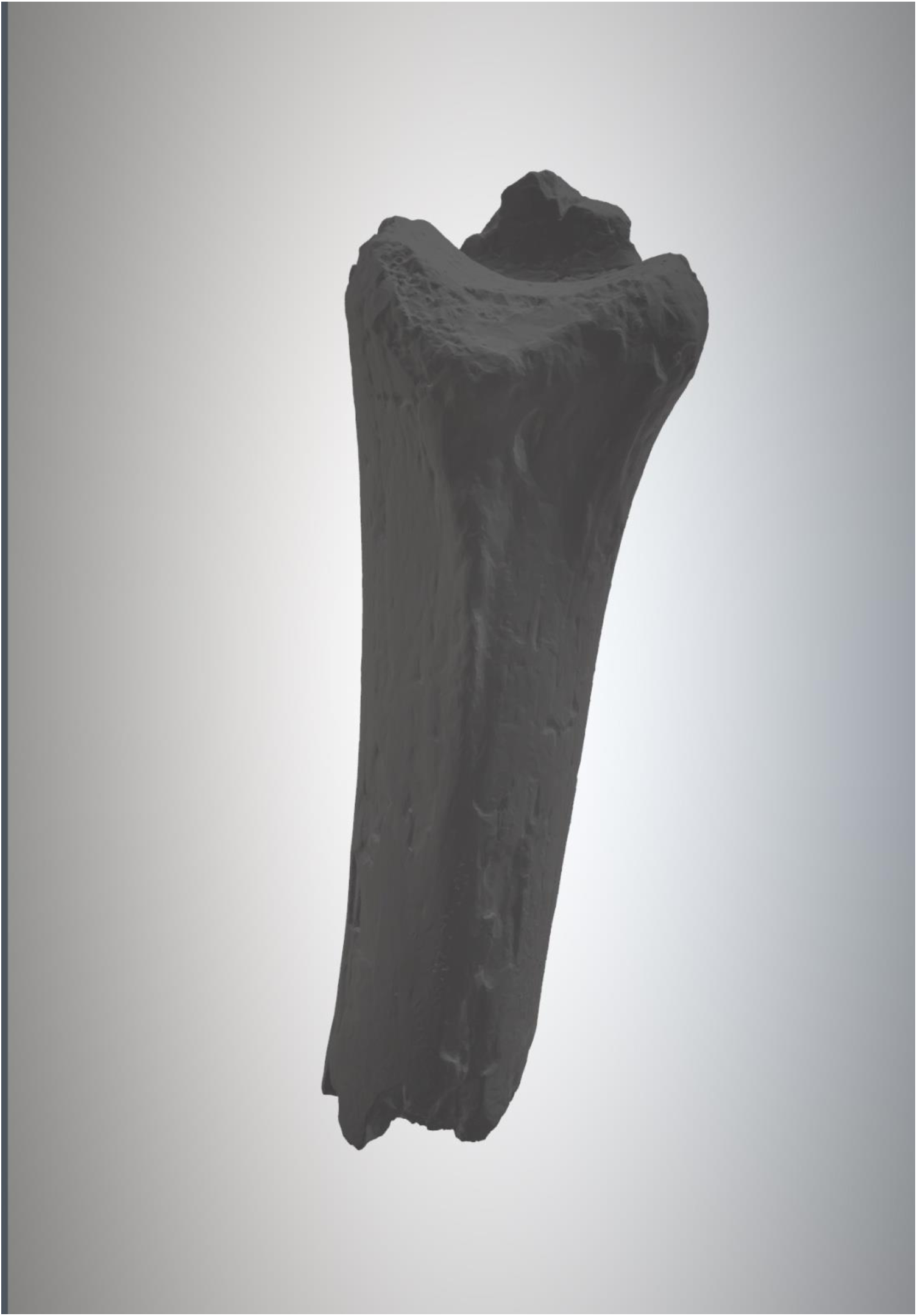
Kanapoi distal right tibia KNMKP 29285 from lateral side, ((downloaded from open source: www.africanfossils.org, Dimensions: x=40.16; y=97.82; z=40.50 mm.)

DeSilva (2009) claimed that the human ankle joint was incapable of dorsiflexion to the extent required for ‘chimpanzee-like’ vertical climbing, and this view has been widely taken on board, particularly by Lovejoy et al. (e.g., 2016). However, Venkataraman et al. (2013a) showed that Twa hunter-gatherers can indeed achieve high ankle dorsiflexion, and engage in vertical climbing since they tend to have longer fibers in the gastrocnemius muscle than neighbouring, non-climbing agricultural communities. The latter is an excellent example of the importance of plasticity - the ability to adapt musculoskeletal anatomy during development to enhance function in the realized niche - to all great apes, including humans, to which we shall return later.

StW 573, KNM-WT15000 and Kanapoi *A. anamensis* thus appear to have very similar proximal and distal tibial morphology, which strongly suggests similarity in function. However, isolated and species-unidentified specimens from Sterkfontein Member 4 often show rather variable morphology. The Member 4 specimen StW 514 assumed to be *A. africanus* by Berger and Tobias (1996), however, combines an *A. anamensis-*like distal tibial condyle (StW 514b) with a proximal condyle (StW 514a), which Berger and Tobias (1996) claimed had distinctly more convex condyles than *A. afarensis.* Organ and Ward (2006), however, found no difference in lateral tibial condyle geometry between StW 514a and *A. afarensis*, and the debate concerning whether, and to what extent, Member 4 australopiths developed a wider range of locomotor adaptations continues. The case of the peculiar pectoral girdle adaptations of *A. sediba* from Malapa, for example (Churchill et al., 2013) is strong evidence that some South African species may have adopted unique modes of postural feeding.

### 3.5. Limb proportions

Figure 23 shows the long bones of the upper and lower limbs of StW 573 compared. At a likely 130 cm tall (RJC pers. comm. to RHC), she was some 10 cm shorter than the average for modern Bolivian women, but some 23 cm taller than AL-288-1 Lucy (106.68 cm according to Jungers, 1988). She was a little shorter than *A. afarensis* KSD-VP-1/1, by the margin which might be expected in a female. Her left humerus’ maximum length is 29 cm; her radius is 24.4 cm long (RJC pers. comm. to RHC), almost identical to the length of the *A. anamensis* radius from Allia Bay, East Turkana, KNM-ER 20419, which it also resembles closely in its (conservative) morphology. Her ulna was 26.3 cm. long. Her total arm length (humerus plus radius) was 53.4 cm. Her femora would have been 33 cm in length, 28.5 for the tibia, giving a total leg length of 61.5 cm (RJC, pers. comm. to RHC and see Heaton *et al*., 2018 submitted). Thus it is no longer a subject for debate whether some early hominids, living at about the time the Laetoli G and S trails were laid down, had hindlimbs that were as long or longer than their forelimbs. StW 573 is the first hominin fossil in which this is unequivocal.

**Figure 23.**
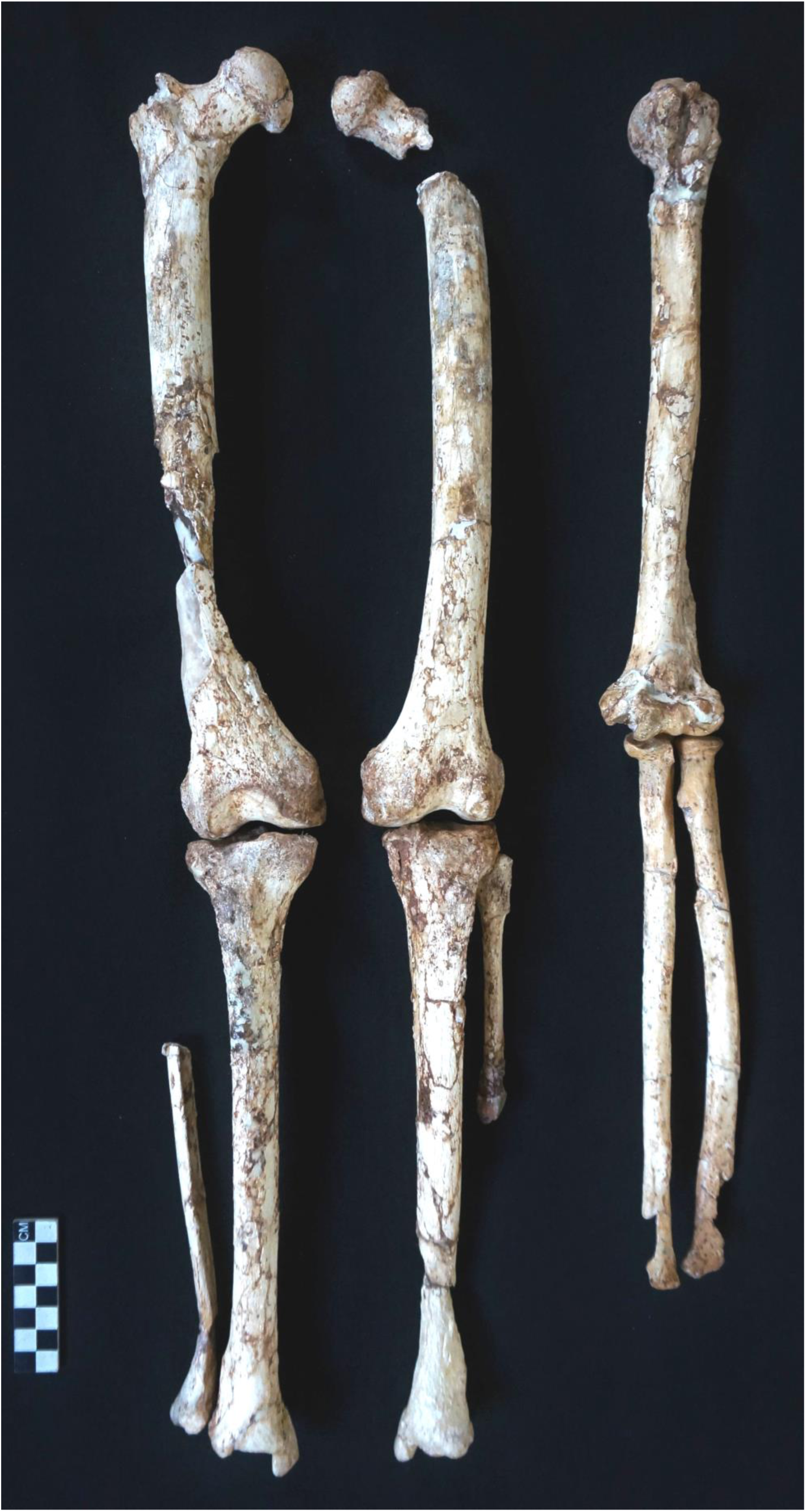
Long bones of the upper and lower limbs of StW 573

**Figure 24.**
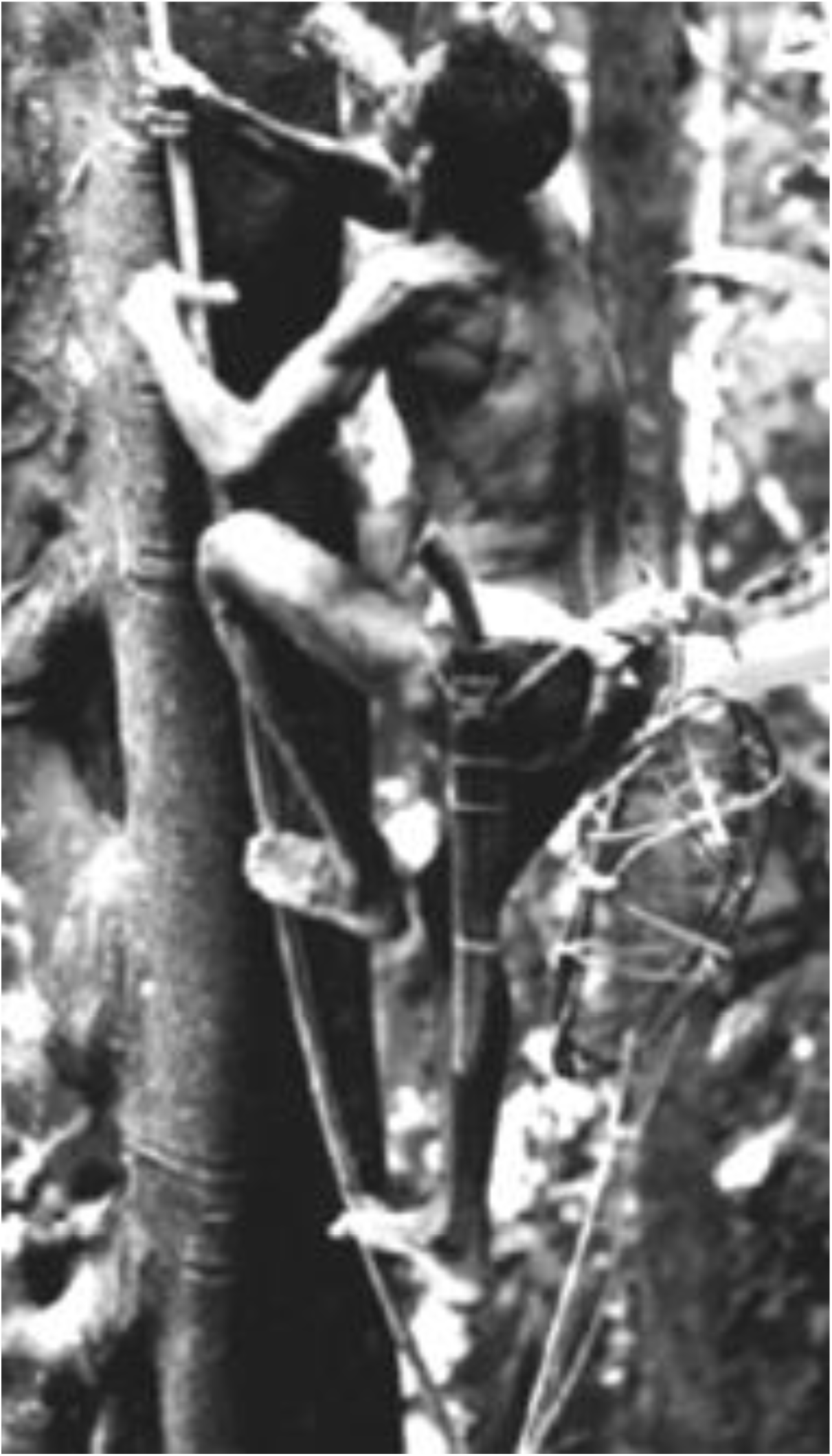
Archival image of an indigenous arboreal forager climbing a thin vine using flexed elbow postures and hallucal grasp (courtesy of Kirk Endicott)

These values give an intermembral index of 86.8 (ratio of h + r L to f + t). Other indices are discussed in Heaton *et al.* (2018 submitted). This is outside and above the human range as reported by Schultz (1937) at 64.5-78, but below that of *Gorilla* at 110-125, that of *Pongo* at 135-150.9, and that of *Pan* at 100.4-100.5, but clearly much closer to the human range than that of the other great apes. The range in *Pan* is so narrow compared to all other great apes as to suggest it is under strong selective control, most likely very tight tuning for effectiveness in quadrupedalism (see Isler *et al*. 2006). Indeed Drapeau and Ward (2007) note that the proportions of the forelimb in *Pan* are highly derived.

## 4. Discussion

### 4.1 Ecomorphology: Testable Hypotheses on Potential Niche

Bock and von Wahlert’s (1965) classic paper, ‘Adaptation and the Form-Function Complex,’ stressed form, function and biological role. It inspired a generation, some to explore biological role by field studies in the natural environment, and others to pursue analyses of the biomechanics of living primates held in captivity. Despite this, it may fairly be said to have had relatively little influence in changing methodology in hominin paleontology, where morphometrics – now most often geometric morphometrics -- continues to dominate research activity, although the introduction of biomechanical modelling techniques such as Finite Elements Stress Analysis and Dynamic Modelling, has been pursued by a (growing) minority. In our view, a newer ecological formulation that is hypothesis- and experiment-driven would be greatly beneficial. This was provided by Wainwright’s (1991) ‘Ecomorphology: Experimental Functional Anatomy for Ecological Problems’. It updates Bock and Von Wahlert (1965) in its focus on performance, and specifically performance of the individual, which is vital because it is the reproductive success of the individual which drives adaptation at population and species levels. Wainwright (1991, p. 680) says: ‘morphology influences ecology by limiting the ability of the individual to perform key tasks in its daily life. In this scheme the effect of morphological variation on behavioral performance is first tested in laboratory experiments. As the behavioral capability of an individual defines the range of ecological resources that it can potentially make use of (the potential niche), the second step in the scheme involves comparing the potential niche of an individual to actual patterns of resource use (the realized niche)’. Those of us who study fossils can rarely carry out ‘laboratory experiments on the effect of morphological variation on behavioural performance’ (p. 680), but increasingly, we can do so in silico, using custom-designed-and-written software. This is usually open-source, such as OpenSim (http://opensim.stanford.edu/work/index.html) and co-author Sellers’ GaitSym (www.animalsimulation.org). The latter has been specifically written for comparative, not human biomechanics, and for palaeontology. Another approach is experimentation using human proxies: we have cited one such study, Johanssen et al. (2017), which tested the effect of light touch on stabilization of the body on unstable supports in a visual simulation of rainforest environments. Similarly, we have just reported that upper limb lengths were short in StW 573 compared to the NHGAs. This suggests less ability to embrace large supports, and particularly, shorter reach, which we hypothesize to reduce the energetic efficiency of arboreal locomotion. Halsey et al. (2017) measured the impact of variation in morphology and locomotor behaviour on the rate of oxygen consumption of 19 elite male parkour athletes as they repeatedly traversed an arboreal-like assault course of 103 m horizontal length. The course consisted of a range of generic gymnasium apparatus such as vaulting horses, raised blocks, high bars, wall bars, and areas filled with loose foam blocks to emulate the range of mechanical conditions present in an arboreal pathway, rather than the exact structure of the forest canopy. Thus, parts of the course incorporated support compliance, irregularity and discontinuity to reflect the conditions experienced during gap crossing between tree crowns, while others were rigid and predicated to reflect the phases between bouts of gap crossing when even large-bodied apes may walk into and out of the core of a tree along thick boughs. They found familiarity with the course had a substantial effect on reducing energetic costs, but there was no evidence to suggest that the locomotor behavior profile of each individual (or the combination of locomotor behaviors that they selected between first and last trials) influenced their ability to attenuate costs. We must therefore, presume more subtle mechanical adjustments are being made to attenuate locomotor challenges. Importantly, athletes with longer arm spans and shorter legs were particularly able to find energetic economies. Thus, our hypothesis that shorter reach would reduce the efficiency of arboreal locomotion is confirmed for one hominin at least, namely *Homo sapiens*. Therefore based on this analogy we conclude that the limb proportions of StW 573 would have reduced her energetic efficiency in arboreal climbing.

A second hypothesis would then be that her long legs and shorter arms would have increased her distance-specific effectiveness in bipedalism. While we have commenced in-silico modelling of StW 573 using sophisticated forwards dynamics modelling under GaitSym, successful fully 3D modelling inevitably takes a great deal of iterative computation, and thus time. However, previous studies of other hominins and of the biomechanical consequences of their body and limb proportions provide strong indications of likely findings. Wang and Crompton (2003, Figure 3), using mass and stature estimates from the literature, found that dimensionless power, mass and stature are closely related, and that humans have arrived at a better combination of these parameters for long distance bipedalism than KNM WT 15000, AL-333 and SK 82. However, as shown by Wang and Crompton (2003, Figure 3) all these fossils occupy a considerably more optimal place on a 3D plot of dimensionless power, mass and stature than for example AL-288-1, OH 62 (a supposed *Homo habilis*—Johanson et al., 1987, but see Clarke, 2017 for a view contra) and Sts 14. Given her estimated stature (130 cm), StW 573 would occupy a position closer to KNM-WT 15000 and AL-333 than to Sts 14, OH 62 and AL-288-1. Thus, our second hypothesis is confirmed by analogy: that StW 573’s distance-specific effectiveness in bipedalism would be enhanced by her longer legs.

On the other hand, following the calculations of Wang et al. (2003), StW 573’s intermembral index of 86.8, outside the human range and larger than that of KNM-WT15000, would not have allowed her to hand-carry loads more than the weight of the upper limb without losing swing symmetry. This contrasts with their estimate that KNM-WT 15000 could effectively carry loads of three times the weight of the upper limb while maintaining swing symmetry. Interestingly, chimpanzees proved unable to hand-carry loads at all without losing swing symmetry, which is interesting in the light of data showing manuports used by chimpanzees in cracking *Panda oleosa* nuts in the Taï forest are carried no more than 10-15 m (Profitt et al., 2018). Similarly, but using inverse dynamics and shoulder-borne loads, Wang and Crompton (2004) showed that, for the given body proportions, KNM-WT 15000 could carry loads of 10-15% body mass for no greater mechanical cost than AL-288-1 would incur walking upright but unloaded. StW 573 would, we predict, function better in this regard than AL-288-1, but by no means as well as KNM-WT 15000. This strongly suggests that her performance capabilities balanced distance-specific terrestrial effectiveness against retention of efficiency in arboreal climbing.

These hypotheses need to be tested, and currently are being tested for StW 573, using forwards dynamic modelling. Again, we can use this technique to discover what advantage would have been delivered to StW 573 by her short femoral neck, given her substantial pelvic flare. Further, as suggested above, given her more cranially oriented glenoid fossa and scapular form (similar to that of *Gorilla* as well as *A. sediba* and KSD-VP-1/1), but her long clavicles (very unlike *A. sediba*) we can use dynamic modelling of her own unique pectoral girdle and pectoral limb architecture to explore the power that she could exert in moderately elevated glenohumeral postures. Thus we will assess the hypothesis that her large brachioradialis flange (suggesting a semiflexed/semipronated elbow posture) would maximize flexor power. This would facilitate climbing on narrow diameter treetrunks and vines with similar kinematics to that recorded for modern human indigenous arboreal foragers, particularly when using hallucal grasping.

### 4.1. Implications of ecological dynamics for functional capabilities of hands and feet

Ecological dynamics seeks to explain coordination and control processes in movement systems during performance of complex multi-articular tasks. This is ner more obvious than during grasping and stepping where multiple rows of bones each form multiple joints controlled by many ligaments and tendons (see Seifert et al., 2016). Here, the hands and feet are interacting directly with the environment and with technology.

The ability of anatomically and morphologically complex organs to adapt efficiently to changes in the environment is driven by the evolutionary mechanism of neurobiological degeneracy. This is the ability of biological elements that are structurally different to perform similar functional outputs (Edelman, 1987). It is quite different to the common engineering concept of redundancy, which refers to the duplication or repetition of similar shaped elements to provide alternative functional outputs in times of mechanical failure (Bernstein, 1967). Therefore multiple means of achieving the same or different functions (according to ecological context) exist by recruitment of structurally different elements. Neurobiological degeneracy ‘is a prominent property of gene networks, neural networks, and evolution itself’ (Edelman and Gally, 2001, p. 13763).

Further, euhominoids/crown hominoids (i.e., Hominidea excluding *eg.* Proconsulidae, Pliopithecidae etc.) display high levels of plasticity in muscle architecture. Venkataraman et al. (2013) showed that (presumably developmental) fibre-length plasticity enables some human forest hunter-gatherers to dorsiflex the ankle to the extent required for chimpanzee-like vertical climbing, while neighbouring non-climbing populations cannot. Further, Neufuss et al. (2014) showed that while lemurs, like all primarily pronograde mammals studied to date, exhibit a dichotomy in axial musculature between deep slow contracting local stabilizer muscles and superficial fast contracting global mobilizers and stabilizers, hominoids, as previously shown for *Homo*, show no regionalization. Thus, it appears that hominoids have been under selective pressure to develop and sustain high functional versatility of the axial musculature, reflecting a wide range of mechanical demands on the trunk in orthogrady. Neufuss et al. (2014) suggest that this is a derived characteristic acquired by early euhominoids. Most likely, this characteristic was acquired by euhominoids such as *Morotopithecus*, or at least *Pierolapithecus*.

Thus, locomotor flexibility is a characteristic of the euhominoid/crown hominoid clade. But in individuals, degeneracy not only stabilizes under perturbation as in light touch (see eg. Johanssen et al. 2017), but helps individuals exhibit adaptability. Multiple alternative recruitment patterns exist in the motor control system, and are variously selected by the CNS (central nervous system) in each grasp or step, as the CNS seeks to optimize performance. It results in functional intra-individual movement variability (Seifert et al., 2014). Thus, it is unsurprising that high intra- and inter-subject variability in human foot pressure cannot be characterized reliably by less than 400 step trials (McClymont 2016; McClymont and Crompton, submitted ms.). Such variability is a natural product of a degenerate system so that, for example, even in small samples, peak midfoot pressures overlap in human, bonobo and orang-utan populations (Bates et al., 2013). Further, prehensive capabilities of the human foot need to be assessed in the context of the greater abduction of the hallux known for many years to exist in habitual barefoot walkers such as Hoffman’s (1905) indigenous forest foragers (and see D’Août et al., 2009). Indigenous human arboreal gatherers such as the Ba’aka, Twa and Batek have the ability to climb small vines using a hallucal grasp (see eg. Figure 25), as observed by Kraft et al. (2014), and equally that of Western adults with reduced pollical capabilities or no pollex to substitute skilled hallucal grasping. Figure 26 illustrates the refined grasp that can be performed by the hallux of some such individuals. The latter, in particular, is an excellent demonstration of how neurobiological degeneracy allows the foot to perform the many fine locomotor skills we tend to associate with the hand. Figure 27 demonstrates that parkour athletes can perform brachiation on an I-beam (here demonstrating the range of plasticity which exists in human finger capabilities, in performing behaviors we normally associate with gibbons and NHGAs,

**Figure 25:**
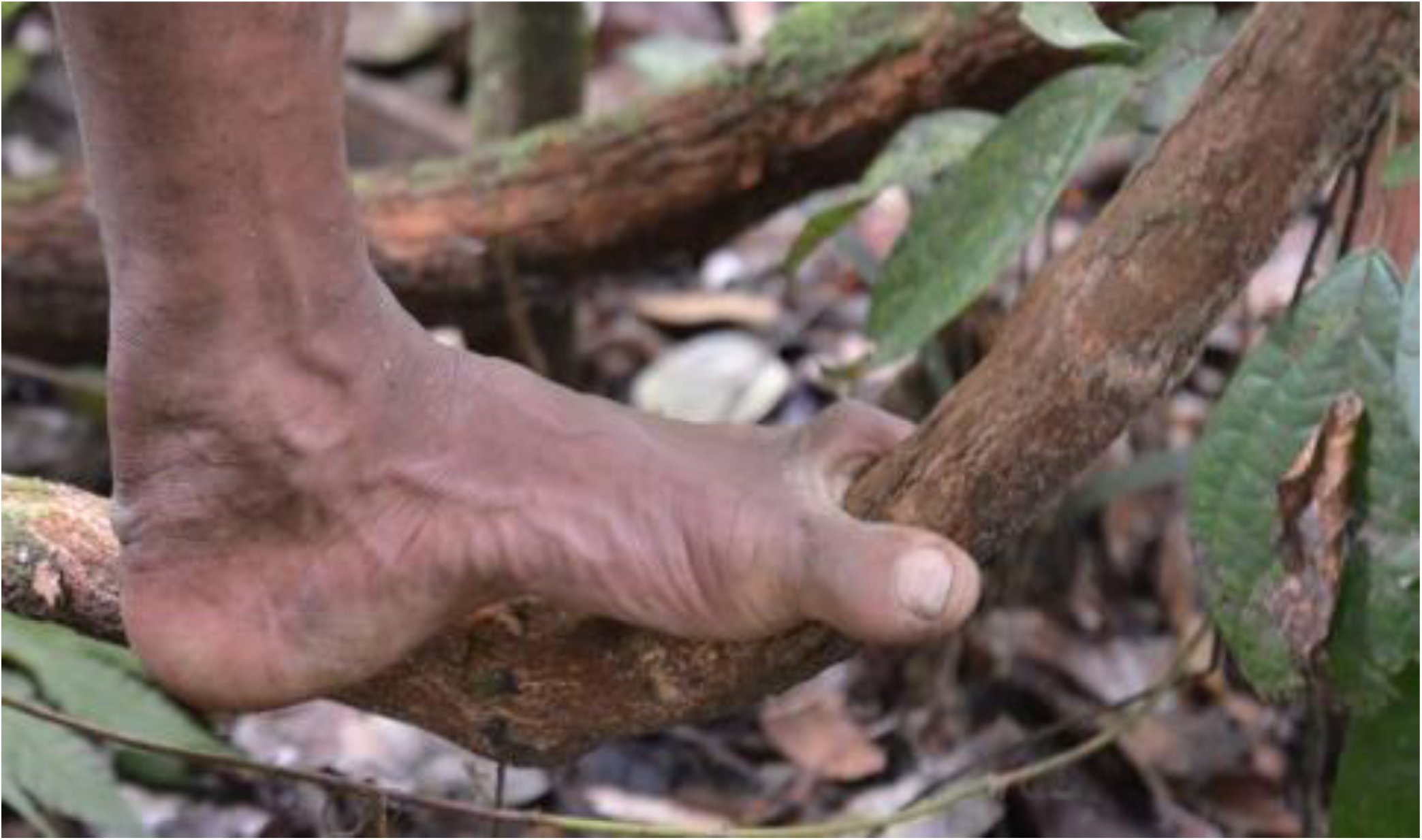
An indigenous Batek arboreal forager demonstrating his hallucal grasp for climbing a small vine (video frame, courtesy of Vivek Venkataraman)

**Figure 26:**
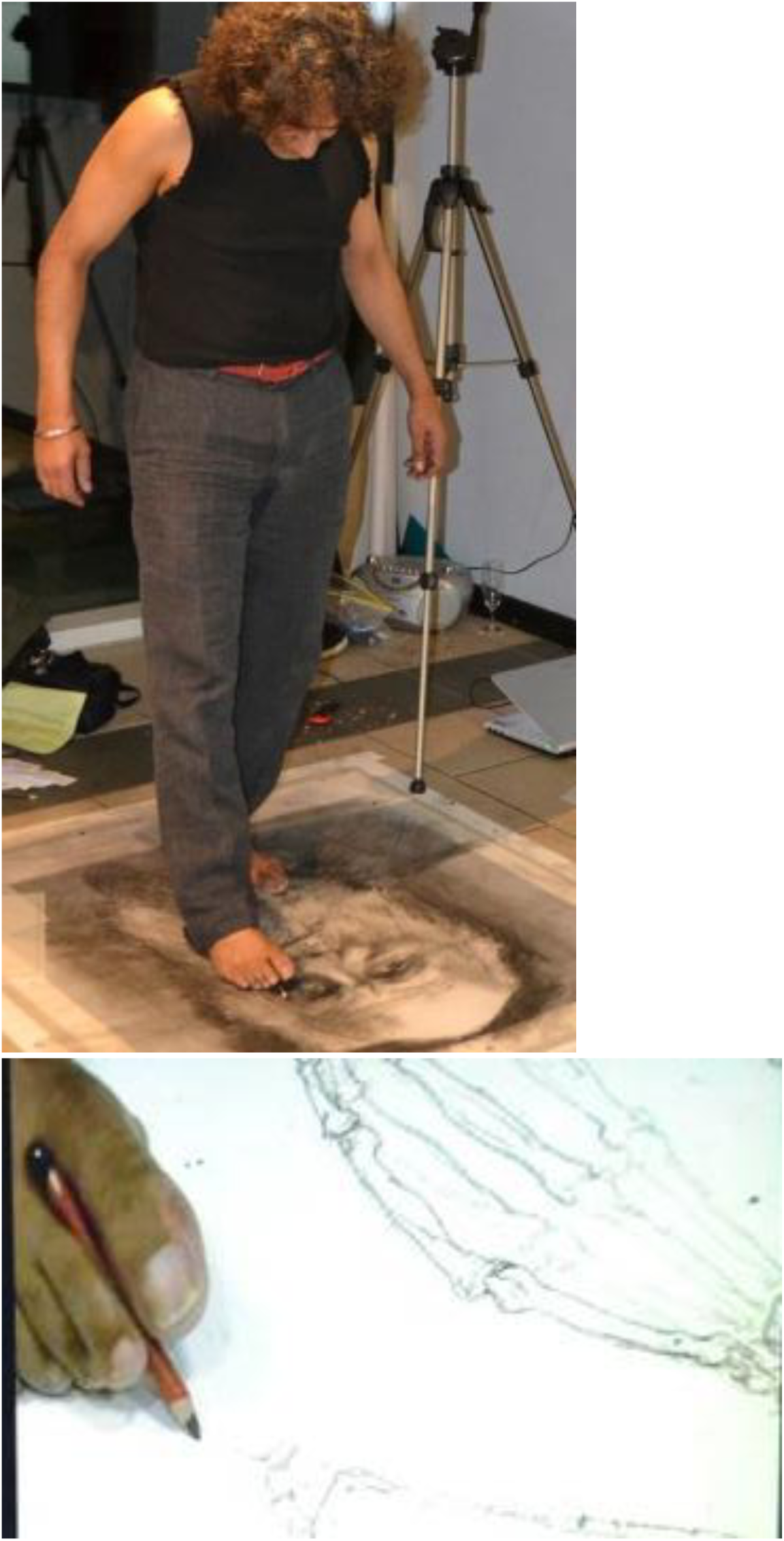
Frames from the official video (’Drawin’ on Darwin’ - Live drawing created at the Meeting of the Primate Society of Great Britain | Saranjit Birdi | Axisweb) of: Top, the artist Saranjit Birdi drawing with his foot Bottom: closeup to show precision of hallucal grasp (courtesy of Emily Saunders and the artist);

**Figure 27:**
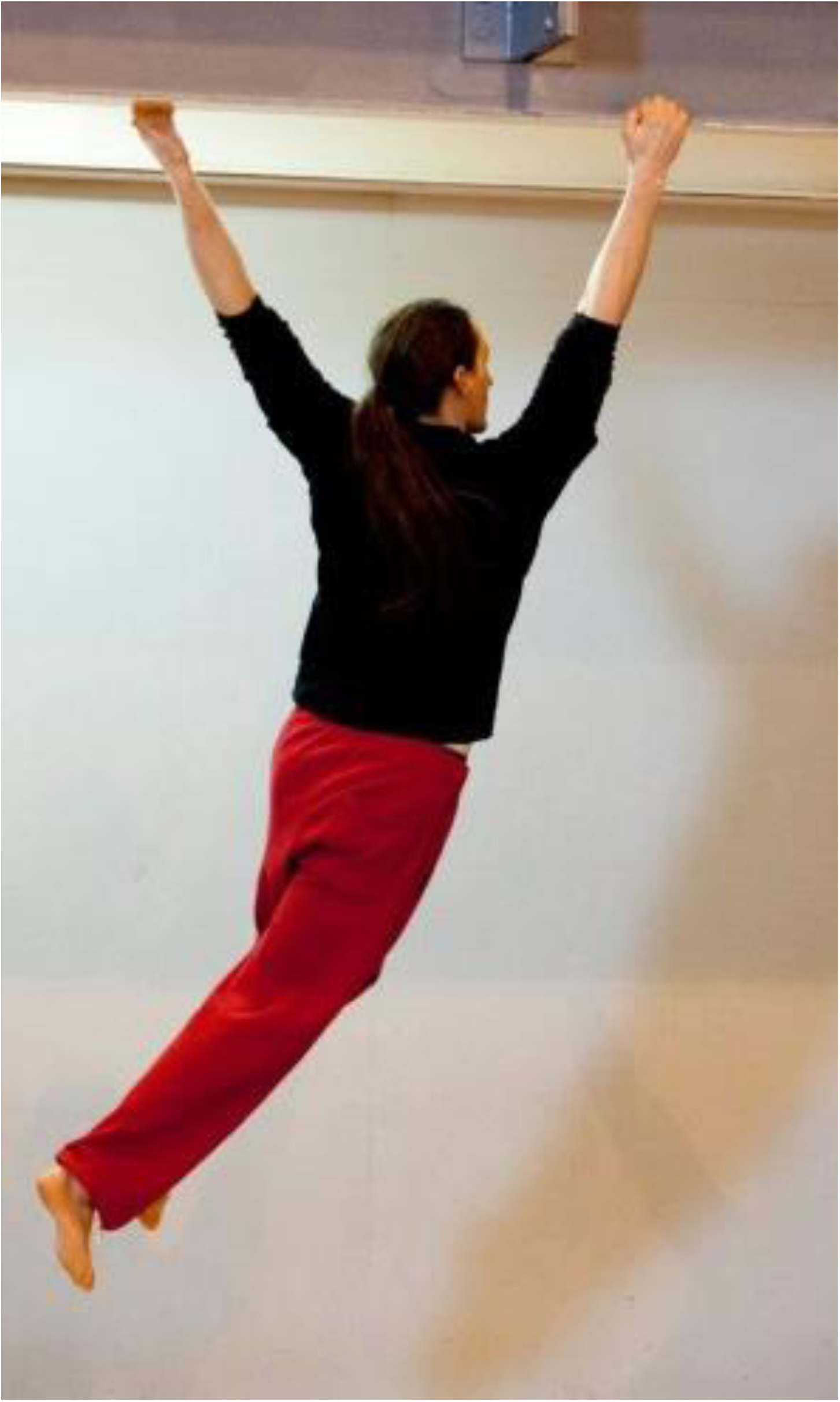
Video frame of a parkour athlete brachiating on an I-beam, demonstrating the plasticity of human finger capabilities (see Halsey et al. 2017, permission of the subject and courtesy Susannah K.S. Thorpe)

The relative proportions of the thumb and fingers of StW 573 (Figure 28) are modern-human-like (Clarke, 1999), as is the case with the *A. afarensis* hand from AL 333 and AL 333w, according to Alba et al. (2003). This suggests that modern human-like hand proportions, as well as grasping capacities (Clarke, 1999, 2002) had their origins in arboreal behaviour before they were exploited in more terrestrial hominins for tool-use. Clarke (2002) notes that no stone tools have been found in Member 2, and there is no suggestion that StW 573 made stone tools. On the contrary, Little Foot’s hand bears a salient apical ridge on the trapezium, a feature commonly present, and marked, in living gorillas (Figure 29). This might have reduced effectiveness of deep, soft opposition (for discussions of prehension see eg. Marzke et al., 1997, and Tocheri et al., 2008).

**Figure 28:**
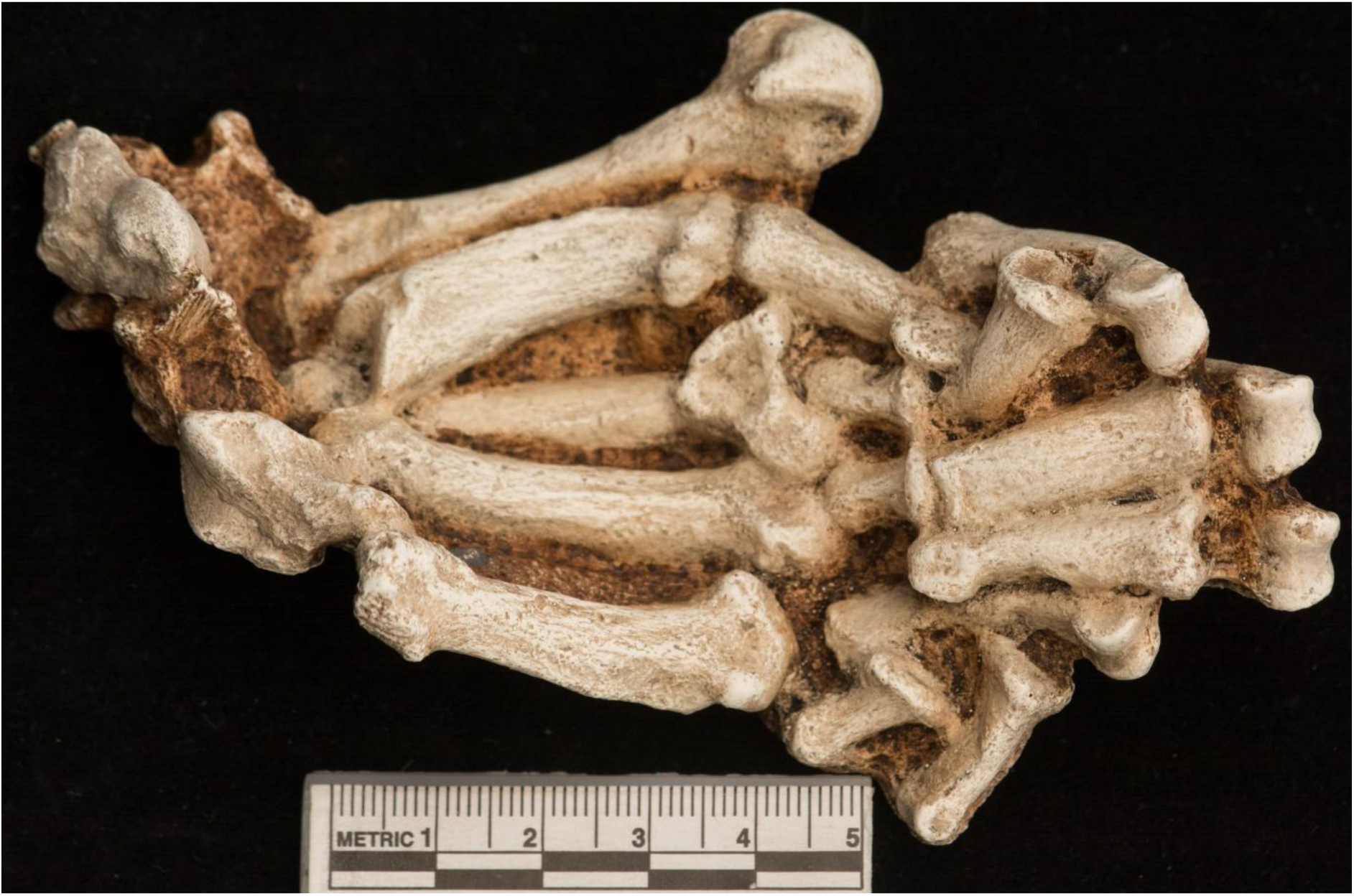
Handbones of StW 573

**Figure 29:**
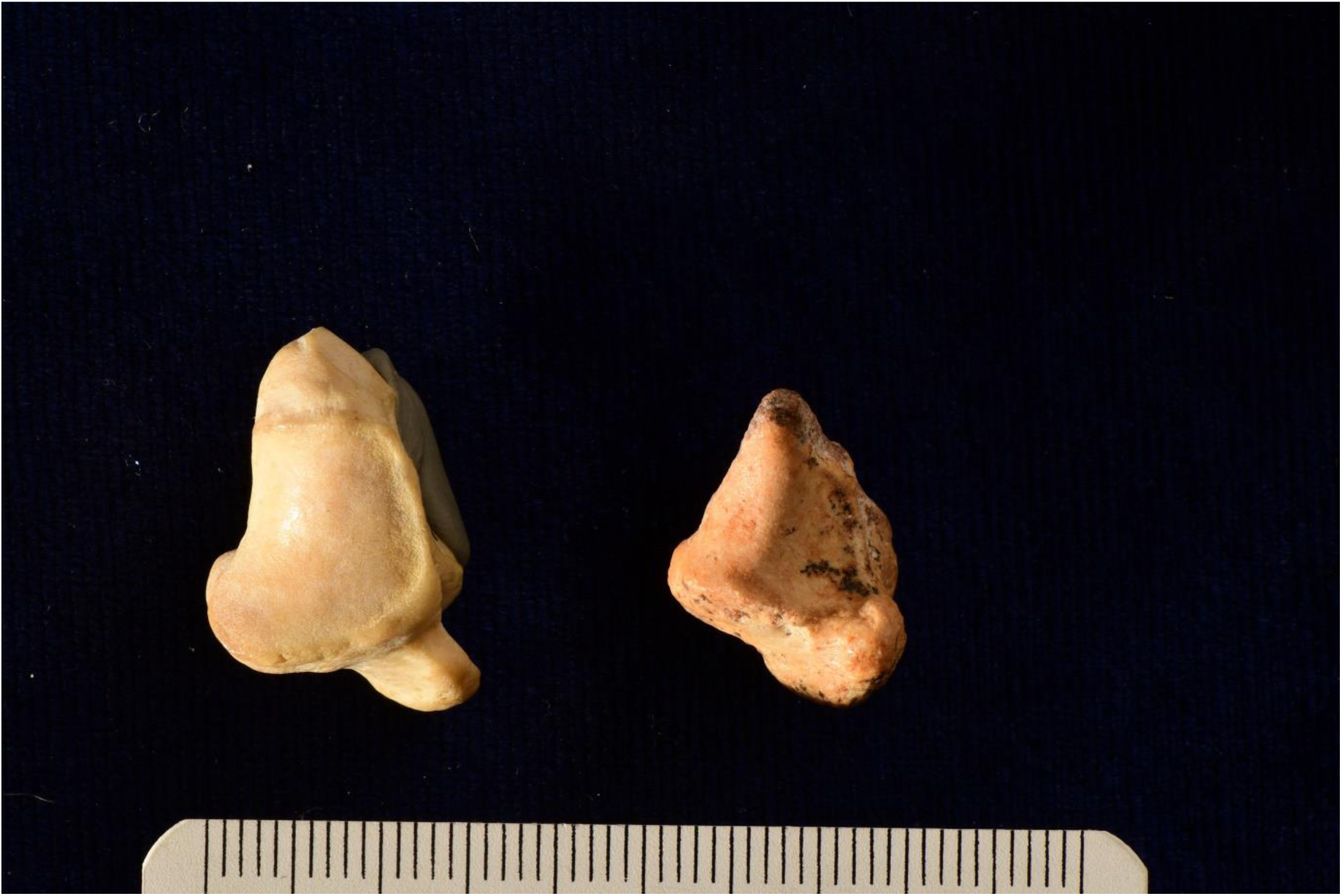
Apical ridge on the trapezium of: left *Gorilla gorilla beringei*, right StW 573 (originals)

Trapezium morphology is highly variable in primates (Napier and Davis, 1959; Hellier and Jeffery, 2006), so care must be taken in interpretation, but it is likely that this structure, absent in humans, might help brace the thumb and its ulnar and radial carpometacarpal and metacarpophalangeal collateral ligaments against forced abduction, similar to ‘gamekeeper’s thumb’ which tends to affect skiers who fall on their hand while still grasping their poles, or football (soccer) goalkeepers who fall while holding a football (Glickel, Barron and Eaton, 1999). In gorillas, the apical ridge might therefore stabilize the pollex in abducted pinch grips during climbing, and we suggest that the case would be the same in StW 573.

Available footbones of StW 573 have been discussed in detail by Clarke (1998, 2002), Clarke and Tobias (1995) and Deloison (2004). Proportions and general morphology broadly resemble those from Woranso-Mille (Haile-Selassie et al., 2012) and Dikika (DeSilva et al., 2018), and the high functional plasticity of the human hallux discussed above must be taken into account in any discussion of hallucal function. Human feet as a whole are highly plastic and functionally degenerate, and as shown by Venkataraman et al. (2013b) and Kraft et al. (2014), they are perfectly capable of functioning efficiently in climbing as well as terrestrial bipedal walking and running, having unquestionably retained a prehensile (if relatively adducted) hallux (see e.g. Figures 25 and 26), contra Holowka and Lieberman (2018). The high human death rates from falls from trees of less than 20 m. quoted by Venkataraman et al. (2013b) are a clear indication that, even were plasticity and degeneracy insufficient, selection would certainly favour retention of hallucal prehension in any human population engaging in barefoot climbing (common in human childhood). It is also highly pertinent to this discussion that analyses of the Laetoli G1 and G2 footprint trails, both of which were formed by hominins penecontemporaneous with StW 573 and KSD-VP-1/1, show that only for very small areas of the foot can external function be statistically distinguished from those made by Holocene human pastoralists and Western humans (McClymont, 2016; McClymont and Crompton submitted ms.). This indicates that the external function of the foot during terrestrial bipedal walking has changed very little since the time of StW 573. Preliminary studies by Raichlen and Gordon (2017) for the new Laetoli S trails are in agreement with this conclusion.

### 4.2 Significance of StW 573 for Hominin origins and the Last Common Ancestor of African Apes

In summaries of the findings in the 2009 special issue of *Science* on *Ardipithecus ramidus*, Lovejoy (p. 74e1) claims ‘*Ar. ramidus* was already well-adapted to bipedality, even though it retained arboreal capabilities. Its postcranial anatomy reveals that locomotion in the chimpanzee/human last common ancestor (hereafter the CLCA) must have retained generalized above-branch quadrupedality, never relying sufficiently on suspension, vertical climbing, or knuckle walking to have elicited any musculoskeletal adaptations to these behaviors.’ While we agree strongly with Lovejoy’s (2009) view, expressed elsewhere in the same paper, that the human/chimpanzee ancestor was not chimpanzee-like, at least in postcranial morphology, we differ with his conclusion that the *Pan/Homo* LCA must have retained ‘generalized above-branch quadrupedality’. *Pan*’s forelimb morphology is highly derived (Drapeau and Ward, 2007) and its intermembral index optimized for quadrupedalism (Isler et al., 2006). Why should the LCA not have been a ‘well-adapted’ arboreal biped as some of us (eg. Crompton et al. 2010) have suggested from field data on *Pongo* and *Gorilla* locomotion? The work of Johannsen et al. (2017) demonstrates clearly that humans retain neural mechanisms for fast response to perturbation in bipedalism on narrow, unstable supports via light touch with the fingers. These would be completely incompatible with a hand loaded in quadrupedal posture.

The skeletal similarity of StW 573 to KSD-VP 1/1 and particularly *A. anamensis*, and evidence for a similar diet to the latter -- substantially C3 foods -- suggests that these hominins had a similar potential niche. It further suggests that the contemporaneous Laetoli G and S trails were made by a very similar hominin which combined continued, if uniquely hominin, modes of arboreal foraging -- in mesic environments -- with effective terrestrial bipedalism. While Ward et al. (2001) concluded that *A. anamensis* was very largely terrestrial, they made a point of not ruling out a substantial arboreal component in its ecology. The postcranial evidence shows that selection was operating on *A. prometheus* to retain considerable arboreal competence: from limb proportions, through the long radius shared with *A. anamensis,* (as indicated by the KNM-ER 20419 Sibilot radius from Allia Bay [see Ward et al. 2001]) to the apical ridge on the trapezium. Indeed the retention of an inner-ear mechanism suited for motion in a complex, 3D environment demonstrated by Beaudet et al. (2018b, submitted) is clear endorsement of the interpretation of a substantially arboreal habitus for *Au. prometheus*. We are thus now able to confirm that the apparent ‘arboreal’ features of early hominins were indeed the subject of positive selection, not selectively neutral anachronisms (see Ward 2002, 2013).

Frequent skeletal similarities of the StW 573 postcranium (e.g., the scapula) to *Gorilla gorilla*, lacking in *Pan*, suggest availability of a similar potential niche, but with reduced use, compared to *Gorilla,* of large tree trunks and increased use of vines and small treetrunks, as noted by Venkataraman *et al.* (2013b) for living human arboreal foragers. Thus we differ also with White et al. (2009, p. 64) in their scenario, which places *Gorilla* on an ‘adaptive pedestal’ separated from australopiths by the chimpanzees, which suggests unidirectional evolution of hominin locomotion. *Pan* is biomechanically highly derived. It is clear that effective arboreal, as well as terrestrial, foraging, albeit less effective than in NHGAs due to adaptations for increased terrestrial effectiveness, were part of the australopith niche and, given locomotor plasticity and degeneracy, remain part of the potential niche of *Homo sapiens* (Kraft et al., 2014).

## 5. Conclusions

Following Wainwright’s (1991) formulation of ecomorphology, we predict that StW 573’s potential niche was exploitation of both arboreal and terrestrial resources, facilitated by plasticity and degeneracy. Toothwear and postcranial similarities to *A. anamensis* suggest a similar primarily C3 diet in mesic mixed forest/grassland. This might include fibrous tubers on the ground and at water margins, as well as tough-skinned arboreal fruit. StW 573 was an effective arboreal biped and climber which had, however, sacrificed some arboreal effectiveness in favour of enhanced energetic efficiency in walking medium to long distances on the ground. She would not have been as effective when load carrying, unlike *Homo ergaster*. Her locomotor posture was upright bipedalism, whether on the ground or on branches, and she was able to stand upright without much muscular activity because of a ‘locking’ or ‘screw-home’ mechanism in the knee which does not seem to have been present in *Ar. ramidus. A. anamensis* and KSD-VP-1/1 probably shared a similar niche. However, we require new fieldwork on lowland gorilla arboreality to establish how the realized niche of *A. prometheus*, *A. anamensis* and *Ar. ramidus* in arboreal foraging might have differed from that of *Gorilla,* accompanied by in-silico testing of locomotor hypotheses concerning early hominin performance capabilities.

## Acknowledgements

This paper was written under an Emeritus Fellowship EM-2017-010 from The Leverhulme Trust to RHC, whose wider research in hominin biology has been primarily funded by The UK Natural Environment Research Foundation and The Leverhulme Trust.

Major funding for the Sterkfontein excavations and MicroCT scanning work have been provided by National Research Foundation grants to KK (#82591 and 82611) and to DS (#98808) and by the Palaeontological Scientific Trust (PAST), without whose support this research would not have been able to continue. We particularly thank Andrea Leenen and Rob Blumenschine for their help in securing major corporate funding, including sustained support from Standard Bank and JP Morgan.

RHC thanks Matt Lotter, Matt Caruana and Kristiaan D’Aout for photographic assistance, and Sarah Elton for her kind advice and support of this special issue. Scans of *Australopithecus anamensis* come from open source: www.africanfossils.org. The authors have no competing interests to declare.

**RHC and JMC dedicate their contribution to the memory of their close friend and colleague, Russ Savage.**

